# Uracil-DNA glycosylase deficiency is associated with repressed tumor cell-intrinsic inflammatory signaling and altered sensitivity to exogenous interferons

**DOI:** 10.1101/2025.07.28.666960

**Authors:** Frank P. Vendetti, Pinakin Pandya, Carina R. Sclafani, Reyna E. Jones, Daniel Ivanov, Robert W. Sobol, Christopher J. Bakkenist

## Abstract

2’-deoxyuridine (dU) is a common DNA lesion resulting from cytosine deamination and from dUMP incorporation by DNA polymerases, both of which are prevalent in cancer. The primary mechanism that repairs dU lesions in genomic DNA is base excision repair initiated by Uracil-DNA Glycosylase 1 (UNG1). We generated *Ung* knockout mouse B16 melanoma cells to investigate the consequences of UNG deficiency in a well-characterized, immunoproficient, syngeneic mouse cancer model. We show that UNG-deficient (ΔUNG) B16 tumors have altered growth kinetics *in vivo* and that their delayed growth is T-cell dependent. Immune profiling revealed reduced CD8^+^ T cell infiltration but augmented CD4^+^ Th1 responses in ΔUNG tumors. *In vitro*, ΔUNG tumor cells exhibit strongly suppressed cell-intrinsic type-I interferon, type-II interferon, and inflammatory signaling gene expression signatures as well as altered cytokine and chemokine secretion. *In vivo*, ΔUNG tumors exhibit a modified inflammatory cytokine and chemokine milieu. Furthermore, ΔUNG tumor cells have altered sensitivity to exogenous interferons *in vitro*, with increased sensitivity to IFN-γ but decreased sensitivity to IFN-α/β. Collectively, our data show that tumor cell-specific UNG deficiency results in an altered tumor microenvironment *in vivo* and provide proof-of-concept data for the use of UNG inhibitors to modulate inflammatory pathways in tumors.

## INTRODUCTION

2’-deoxyuridine (dU) is a common DNA lesion resulting from cytosine deamination and from dUMP incorporation by DNA polymerases, both of which are prevalent in cancer (1). Many cancers harbor activating mutations in cytidine deaminases, driving widespread dU lesions, with cytidine deamination representing a major mutational signature in human cancers (2–4). Additionally, many cancer patients receive antimetabolites like 5-fluorouridine (5-FU) and pemetrexed, which increase the dUTP/TTP ratio, leading to higher dUMP incorporation into DNA (5). dU lesions in nuclear DNA are excised by uracil-DNA glycosylases UNG1, UNG2, and SMUG1 (6). UNG2 binds PCNA and removes dU from RPA-coated single-stranded DNA at the replisome, while UNG1 and SMUG1 excise dU in non-replicating chromosomal regions (7, 8). In short-patch base excision repair (BER), APE1 hydrolyzes the abasic site, Polβ inserts the correct nucleotide, and Ligase III seals the nick, while in long-patch BER, Polβ or Polε and FEN1 replace eleven nucleotides and Ligase I seals the nick (9).

Antimetabolites can create futile repair cycles, where dU is continuously excised and reinserted by DNA polymerases, allowing dU lesions to persist in chromosomal DNA. Persistent dU lesions have been associated with replication stress, chromosome instability, and the efficacy of cancer therapies *in vitro* and in immune deficient mouse models of cancer (10, 11). Furthermore, persistent dU lesions lead to under-replication of DNA and chromosomal instability in homologous recombination-deficient (HRD) cancer cells *in vitro*, and UNG inhibition or depletion is synthetic lethal with homologous recombination deficiency *in vitro* (12). Thus, the repair of dU lesions in cancer patients treated with antimetabolites and clastogens is likely to contribute to the formation of extrachromosomal DNA fragments and, in some instances, these fragments may contain dU lesions (10, 13). UNG is overexpressed in many human cancers suggesting that the repair of dU lesions is critical for tumorigenesis and UNG and APE1 inhibitors are in various stages of clinical development (14–19). UNG and APE1 inhibitors are likely to increases extrachromosomal DNA fragments containing dU lesions and abasic sites, respectively. Disrupting UNG-dependent repair, by knockdown or deletion of UNG, or by APE1 inhibition, sensitizes human cancer cells to antimetabolites *in vitro* and in immune deficient mouse models of cancer (20–22). However, studies using UNG-deficient tumor models in immunoproficient hosts are lacking, and how the loss of UNG activity in tumors may affect the tumor immune microenvironment is unknown. This is important, as recent preclinical data in mice suggest that antimetabolites of interest for combination with UNG inhibitors, including 5-FU and pemetrexed, are immunomodulatory on their own, and can prime tumors to respond to immunotherapy (23, 24). Here, we use the well-characterized, immunoproficient, syngeneic mouse B16 melanoma model to investigate the consequences of tumor cell-specific UNG enzyme deficiency on the tumor immune microenvironment.

## RESULTS

### UNG deficiency alters B16 melanoma growth in immunocompetent mice and improves responsiveness to anti-PD-L1 therapy

We generated clonal *Ung* knockout B16 cell lines using CRISPR/Cas9, thereby targeting both protein isoforms, UNG1 and UNG2. One clone, named ΔUNG, has a homozygous deletion that generates a frameshift and a stop codon, while a second clone, named ΔUNG_2, has a homozygous deletion that deletes two proline residues **(Supp Fig S1).** As expected, both clones exhibit significantly reduced glycosylase activity at dU:dA and dU:dG base pairs, and ΔUNG exhibits no loss of APE1 activity **(Supp Fig S2)**. Therefore, we used ΔUNG throughout this study.

To determine whether tumor cell-specific UNG deficiency impacts tumor growth *in vivo*, we inoculated immunocompetent C57BL/6 mice with ΔUNG or Cas9-expressing control B16 cells and measured tumor growth over time. ΔUNG tumors exhibited delayed early tumor growth (within the first 2 weeks) but accelerated rapidly at later time points (after day 17) **(Fig 1A, Supp Fig S3A)**. These differences in early tumor growth were significant and reproducible across multiple experiments **(Fig 1B)**. The altered growth of ΔUNG tumors was not due to differences in cell proliferation, as ΔUNG and control cells proliferated at the same rate in *vitro* **(Fig 1C).** Furthermore, ΔUNG and control tumors grew at the same rate in athymic nude mice **(Fig 1D)**, indicating that the altered growth kinetics of ΔUNG tumors in immunocompetent C57BL/6 is dependent on the immune system, in particular, T cells.

**Figure 1.**
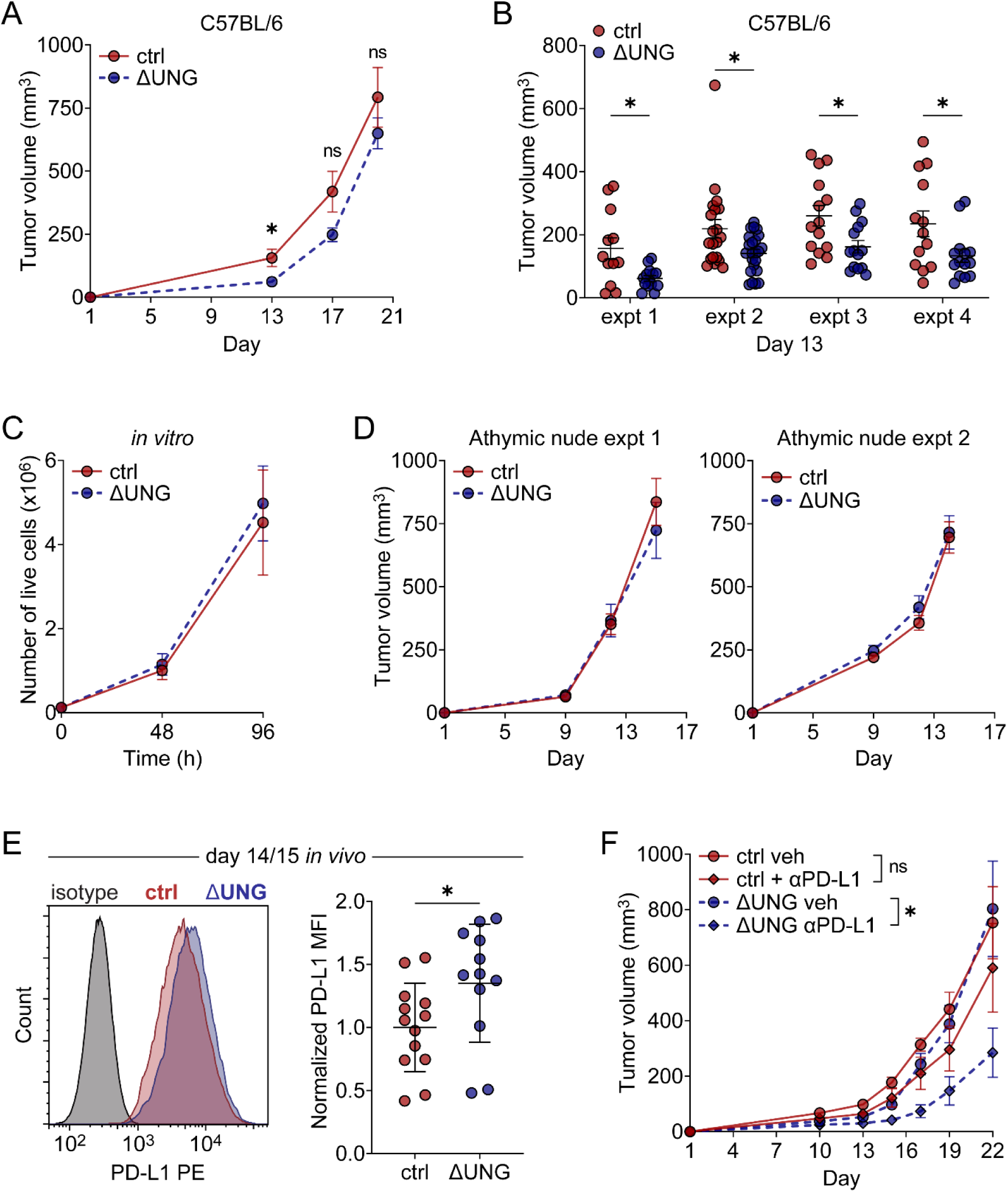
UNG deficiency alters B16 melanoma growth in immunocompetent mice and improves responsiveness to anti-PD-L1 therapy. **A.** Growth of Cas9-expressing control (ctrl) and UNG-deficient (ΔUNG) B16 tumors in immunocompetent C57BL/6 mice. Mean tumor volumes and SEM bars shown. Data from one experiment. n = 12 (ctrl) or 14 (ΔUNG) mice. **B**. Individual ctrl and ΔUNG tumor volumes in C57BL/6 mice on day 13 from four independent experiments. Mean ± SEM bars shown. A-B. *p<0.05, ns (not significant) by two-tailed, unpaired Welch’s t test at the indicated measurement time points. **C.** Growth of ctrl and ΔUNG B16 cells *in vitro* quantified as the number of live cells. Mean and SD bars shown. Data from one experiment with 5 biological replicates. **D.** Growth of ctrl and ΔUNG B16 tumors in immunocompromised athymic nude mice from two independent experiments. Mean tumor volumes and SEM bars shown. n = 19 (expt 1) and n = 18 (expt 2 ctrl) or 19 (expt 2 ΔUNG) mice per group. **E.** Flow cytometry analysis of PD-L1 expression on ctrl and ΔUNG B16 tumors (CD45neg cells) on day 14-15 in C57BL/6 mice. (Left) Histogram showing the median fluorescence intensity (MFI) of PD-L1 PE staining on representative ctrl and ΔUNG B16 tumors, compared to isotype control staining. (Right) PD-L1 MFI on ctrl and ΔUNG B16 tumors normalized to the mean MFI of ctrl control tumors. Data combined from 2 independent experiments, each with 4-8 mice per group, for total n = 13 (WT) or 12 (ΔUNG) mice. Mean ± SD bars shown. *p<0.05 by two-tailed, unpaired Welch’s t test. **F.** Ctrl and ΔUNG B16 cells were injected (day 1) into C57BL/6 mice and mice were treated with 100 µg anti-PDL1 every 3 days for 6 doses, starting on day 2. Mean tumor volumes ± SEM bars shown. *p<0.05, ns (not significant) by mixed-effects analysis of tumor growth data over time with Holm-Sidak adjustment for multiple comparisons.

Since the early delayed growth of ΔUNG tumors was T cell-dependent and was not sustained, we hypothesized that increased PD-L1 expression, and the subsequent suppression of T cell activity, may be responsible for accelerating tumor growth at later time points. We found elevated tumor PD-L1 expression at days 14-15 in ΔUNG compared to control tumors **(Fig 1E)**. We hypothesized that PD-1/PD-L1 blockade may improve the durability of the T cell-dependent growth delay in ΔUNG tumors. The B16 tumor model is known to be relatively resistant to immune checkpoint blockade, either via anti-PD-1 or anti-PD-L1 (25–27). To determine whether UNG deficiency alters the sensitivity of B16 tumors to checkpoint blockade, we treated mice with anti-PD-L1 antibody every 3 days for 6 doses, beginning one day after implantation of ΔUNG or control cells. While control tumors treated with anti-PD-L1 exhibited some growth inhibition compared to vehicle-treated control tumors, the differences were not statistically significant. Conversely, ΔUNG tumor growth was significantly inhibited by anti-PD-L1 therapy compared to growth of vehicle-treated ΔUNG tumors **(Fig 1F and Supp Fig S3B)**. Therefore, our data support that UNG activity in tumor cells plays a role in mediating the tumor-immune microenvironment and responsiveness to immunotherapy.

### UNG deficiency impacts the tumor microenvironment and promotes a CD4^+^ Th1 response

To investigate how T cell responses are reshaped in UNG-deficient tumors, we profiled the immune infiltrates in ΔUNG and control B16 tumors at days 10 and 14 after implantation. At day 10, ΔUNG tumors exhibited significantly increased CD4^+^ T cells (conventional and regulatory), as percentages of the total CD45^+^ immune cell (CD45^+^) infiltrate, compared to control tumors **(Fig 2A**, right panel). While not statistically significant, ΔUNG tumors, on average, also had reduced CD8^+^ T cell infiltration **(Fig 2A**, left panel). There were no differences in overall T cell infiltration (quantified as Thy1.2^+^ cells) **(Fig 2B)** or in total CD45^+^ immune cell infiltration **(Fig 2C)** between control and ΔUNG tumors at day 10. We also observed no differences in the infiltration of NK/NKT cells, B cells, or myeloid-lineage cells in control and ΔUNG tumors at day 10 **(Supp Fig S4A-C)**.

**Figure 2.**
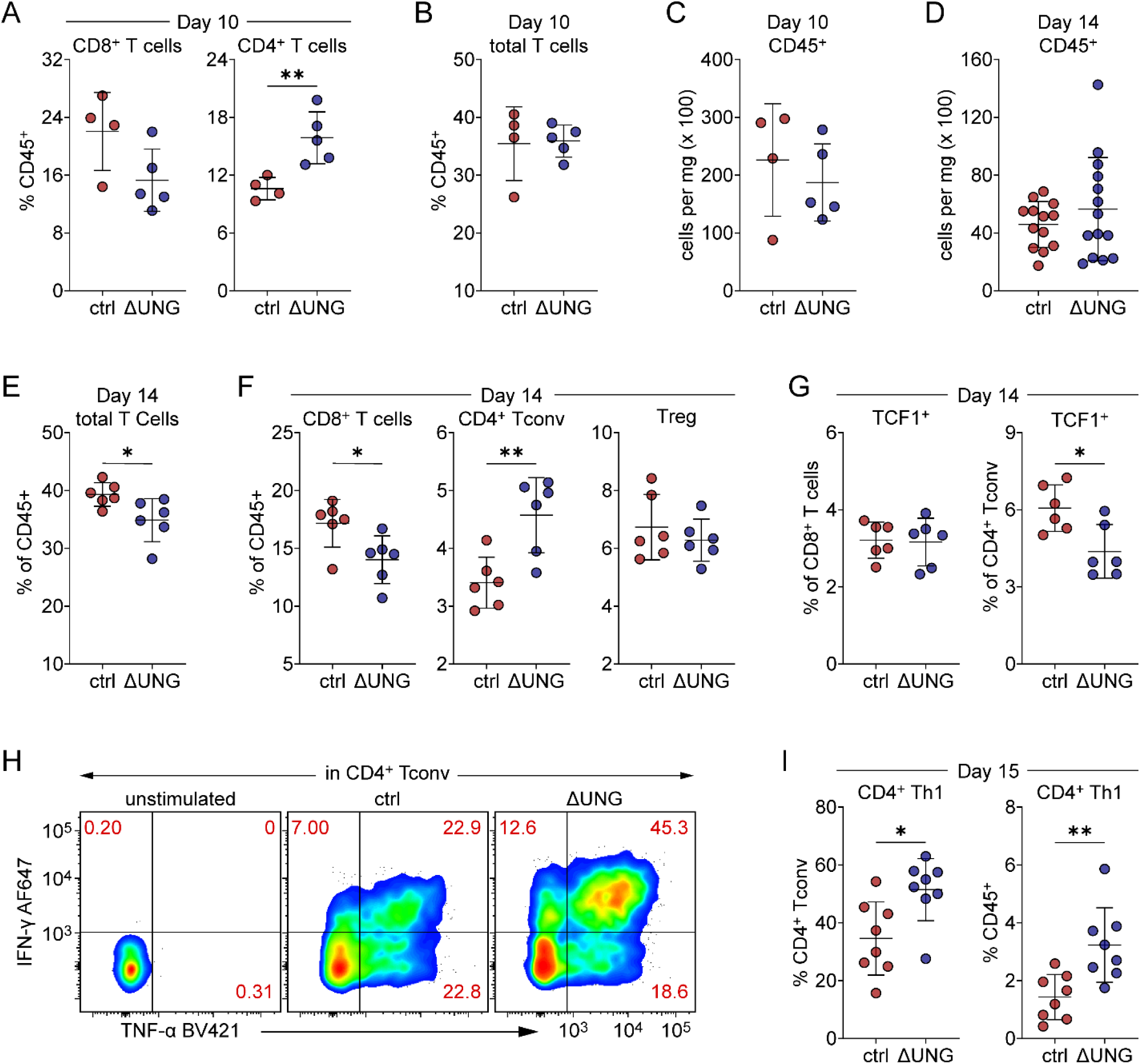
UNG deficiency alters the B16 tumor infiltrate and promotes a CD4^+^ Th1 response. A-C. Flow cytometry immunoprofiling of control (ctrl) and ΔUNG B16 tumors on day 10. Data from one experiment with n = 4 (ctrl) or 5 (ΔUNG) mice per group. **A-B.** Quantitation of tumor-infiltrating T cell populations as percentages of the CD45^+^ immune cell infiltrate. Quantified populations include CD8^+^ and CD4^+^ **(A)** and total **(B)** T cells. **C.** Quantitation of the relative number of tumor-infiltrating CD45^+^ immune cells per mg of tumor stained. **D-G.** Flow cytometry immunoprofiling of ctrl and ΔUNG B16 tumors on day 14. **D.** Quantitation of the relative number of tumor-infiltrating CD45^+^ immune cells per mg of tumor stained. Data combined from 2 independent experiments, each with 6-8 mice per group, for total n = 13 (ctrl) or 14 (ΔUNG) mice. **E-F.** Quantitation of tumor-infiltrating T cell populations as percentages of the CD45^+^ immune cell infiltrate. Quantified populations include total T cells **(E)** and T cell subsets **(F)**: CD8^+^, conventional CD4^+^ (Tconv), and regulatory (Treg) T cells. **G.** Quantitation of TCF1-expressing CD8+ T cells and CD4+ Tconv as percentages of the parent populations. **E-G.** Data from one experiment. n = 6 mice per group. **H-I.** Flow cytometry analysis of IFN-γ and TNF-α production by CD4^+^ Tconv on day 15 following *ex vivo* stimulation of ctrl and ΔUNG tumor infiltrates with PMA/ionomycin. **H.** Cytograms showing IFN-γ and TNF-α staining in representative stimulated ctrl and ΔUNG tumor infiltrates, compared to unstimulated control. **G.** Quantitation of tumor-infiltrating CD4^+^ Th1 cells, defined as IFN-γ^+^ (TNF-α^+/-^) CD4^+^ Tconv, as percentages of parental CD4^+^ Tconv or the CD45^+^ immune cell infiltrate. Data from one experiment. n = 8 mice per group. **A-G, I.** Mean ± SD bars shown. *p<0.05, **p<0.01, ns (not significant) by two-tailed, unpaired Welch’s t test.

Next, we profiled T cell populations at day 14 in one experiment and myeloid cell populations at day 14 in a second experiment. Collectively, from the two experiments, we observed no significant differences in the total CD45^+^ immune cell infiltrate **(Fig 2D)**. However, ΔUNG tumors had significantly decreased total CD3^+^ cells (encompasses all T cells), as percentages of the immune infiltrate, compared to control tumors **(Fig 2E)**. The decrease in total CD3^+^ cells in ΔUNG tumors was associated with significant decreases in CD8^+^ T cells but significant increases in conventional CD4^+^ T cells (CD4^+^ Tconv), with no changes in regulatory T cells (Treg) **(Fig 2F)**. We observed minimal differences in the myeloid populations profiled at day 14 **(Supp Fig S4D-F)**, except for reduced CD103^+^ dendritic cells (DC) in ΔUNG tumors **(Supp Fig S4D)**. As CD103^+^ DC are important for priming the anti-tumor CD8^+^ T cell response (28), the reduction in this antigen-presenting population may explain, at least in part, the reduced CD8^+^ T cells in ΔUNG tumors.

Since we noted increased tumor PD-L1 expression in ΔUNG compared to control tumors *in vivo* **(Fig 1E)**, and the type II-IFN, IFN-γ, is a primary mediator of PD-L1 upregulation by cells *in vivo* (29–31), we examined the cytokine competency of tumor-infiltrating T cells at day 15 to determine whether increased T cell functionality was responsible for elevated PD-L1 in ΔUNG tumors. We examined IFN-γ and TNF-α production by CD4^+^ Tconv **(Fig 2H-I)** and CD8^+^ T cells **(Supp Fig S5A-B)**, and IL-17 production by CD4^+^ Tconv **(Supp Fig S5C)**, following stimulation *ex vivo* with PMA/ionomycin. ΔUNG tumors were significantly enriched in IFN-γ-competent (± TNF-α) CD4^+^ T cells, which are Th1 effector CD4^+^ T cells, as percentages of both the parent CD4^+^ Tconv population and the total CD45^+^ immune infiltrate **(Fig 2I)**. Conversely, we saw no significant differences in IFN-γ/TNF-α-competent CD8^+^ T cells in control versus ΔUNG tumors **(Supp Fig S5A-B)**. Similarly, we saw no significant differences in IL-17-competent CD4^+^ Tconv (i.e., Th17 cells) in control versus ΔUNG tumor-infiltrates **(Supp Fig S5C)**. Since ΔUNG tumors contained similar proportions of cytokine-competent CD8^+^ T cells and fewer CD8^+^ T cells overall compared to control tumors, the enriched CD4^+^ Th1 cell response was a likely source of IFN-γ driven PD-L1 expression in ΔUNG tumors.

### UNG deficiency represses tumor cell-intrinsic basal and ATRi-induced inflammatory signaling

Given that UNG deficiency did not alter B16 cell growth *in vitro* or tumor growth in immunocompromised mice but did impact B16 tumor growth and immune infiltration in immunocompetent mice, we reasoned that UNG deficiency altered pathways responsible for communication with the immune system. To investigate this, we performed RNA sequencing to determine the genes altered in UNG-deficient B16 cells in an unbiased manner. RNA-seq and gene set enrichment analyses (GSEA) revealed that the principal gene sets in which ΔUNG cells are deficient in basal expression are the type-I IFN response (53/89 genes), the type-II IFN response (79/156 genes), and the inflammatory response (50/122 genes) **(Fig 3A).** The second UNG-deficient clone, ΔUNG_2, is deficient in basal expression of these same gene sets: the type-I IFN response (60/80 genes), the type-II IFN response (81/156 genes), and the inflammatory response (67/122 genes) **(Fig 3A)**. Gene ontology (GO) analyses yielded multiple biological pathways, including viral responses, innate and effector immune responses, and interferon-beta response, that are altered in ΔUNG versus control cells, and that are consistent with the principal gene sets identified by GSEA **(Supp Fig S6A)**.

**Figure 3.**
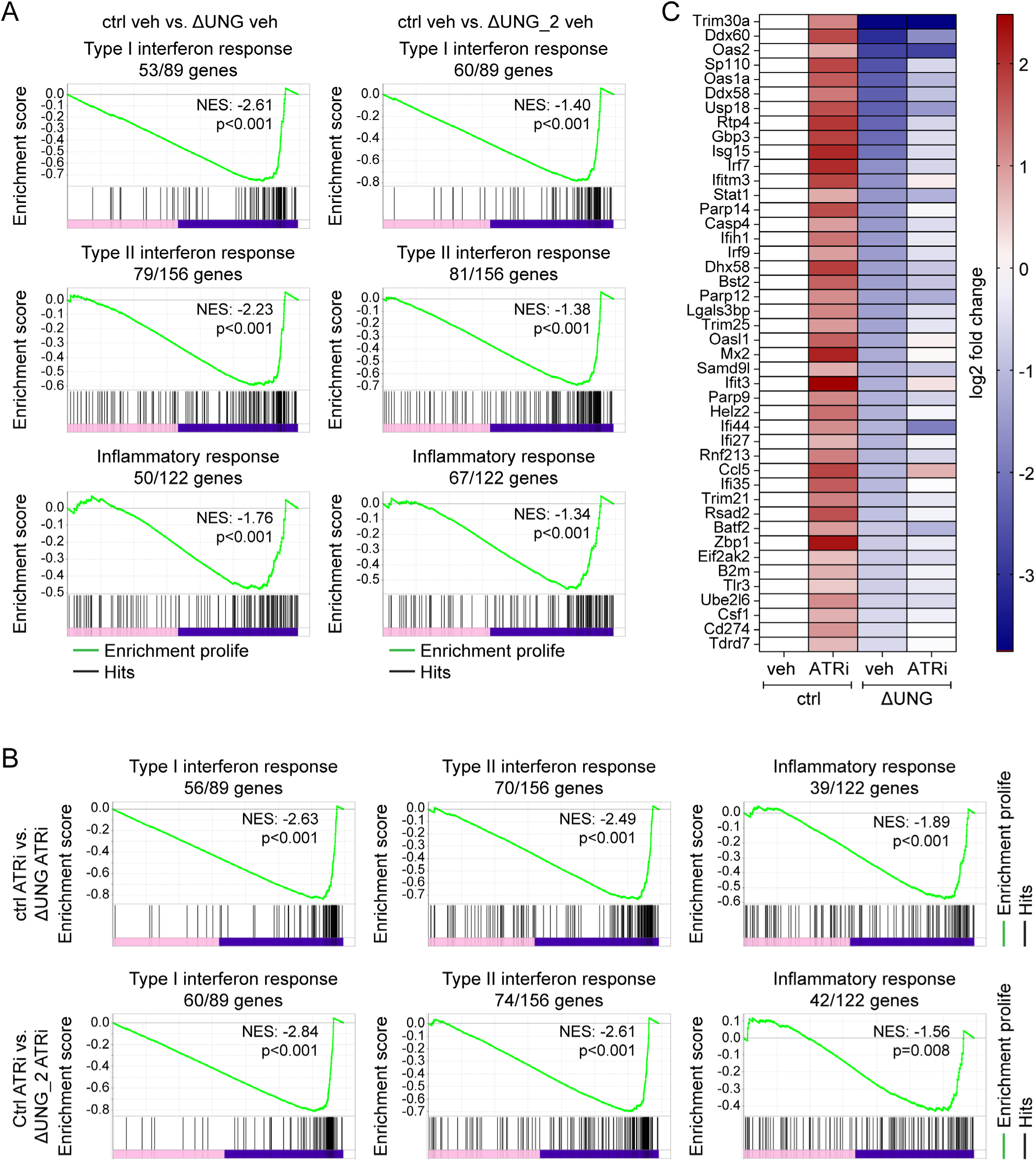
UNG deficiency represses tumor cell-intrinsic basal and ATRi-induced inflammatory signaling. A-C. Control (ctrl) and two clones of UNG-deficient (ΔUNG and ΔUNG_2) B16 cells were treated with vehicle (veh) or 5 µM AZD6738 (ATRi) for 48 h. RNA sequencing was performed with two biological replicates per line and condition (one replicate for ATRi-treated ΔUNG_2). **A-B.** Gene set enrichment analysis (GSEA) of the most significantly changed pathways in veh-treated **(A)** and ATRi-treated **(B)** cells. NES = normalized enrichment score. **C.** Heatmap depicting the most significantly changed genes associated with the inflammatory signaling pathways identified by GSEA.

ATR inhibitors (ATRi) potentiate radiation-induced inflammation in patients and mouse tumor models and induce the type-I interferon (IFN-α/IFN-β) signaling in cells (32–36). Therefore, we performed RNA-seq on ATRi-treated (5 µM AZD6738) control, ΔUNG, and ΔUNG_2 B16 cells to determine whether UNG deficiency attenuates ATRi-induced inflammatory signaling pathways. GSEA revealed that the principal gene sets in which ΔUNG and ΔUNG_2 cells are deficient in ATRi-induced expression are the same gene sets (type-I IFN response, type-II IFN response, inflammatory response) repressed at baseline in ΔUNG and ΔUNG_2 **(Fig 3B).** GO analyses again yielded multiple biological pathways (viral responses, interferon-beta response) that are altered in ATRi-treated ΔUNG versus ATRi-treated control cells, and that are consistent with the principal gene sets identified by GSEA **(Supp Fig S6B)**. A heat map of the top altered genes identified by GSEA demonstrates that multiple interferon-stimulated genes and inflammatory genes are repressed in basal expression in ΔUNG versus control cells **(Fig 3C)**. While ATRi induces the expression of many of these genes in ΔUNG cells, their induction is markedly attenuated when compared to ATRi-induced expression of these genes in control cells **(Fig 3C)**. Broadly speaking, UNG deficiency represses both basal and ATRi-induced expression of genes associated with interferon-stimulated, inflammatory, and innate immune signaling pathways.

### UNG deficiency alters the cytokine and chemokine secretion by tumor cells *in vitro*

Next, we investigated whether the gene expression changes identified by RNA-seq equated to changes in the secretion of inflammatory cytokines and chemokines from tumor cells *in vitro*. We performed secretome analyses of supernatants from vehicle-treated and ATRi-treated control, ΔUNG, and ΔUNG_2 B16 tumor cells at 48 h using the mouse Cytokine/Chemokine 44-Plex Discovery Assay Array (Eve Technologies). Except for VEGF and Fractalkine, the majority of basal cytokine and chemokine secretion was reduced or unchanged in ΔUNG and ΔUNG_2 cells compared to control cells **(Fig 4A-B)**. This was most striking for inflammatory cytokines, including IL-16 and the IL-6 family (IL-6, IL-11, LIF), and the inflammatory chemokine KC **(Fig 4A-B)**. Secretion of multiple inflammatory cytokines (IL-6, IL-11, and LIF) and chemokines (IP-10 and MCP-5) from ATRi-treated ΔUNG and ΔUNG_2 cells was reduced compared to ATRi-induced secretion from control cells **(Fig 4C-D)**. In several cases, secretion of inflammatory cytokines (IFN-β and IL-16) and chemokines (KC and RANTES) from ATRi-treated ΔUNG and ΔUNG_2 cells was reduced compared to even basal secretion from vehicle-treated control cells **(Fig 4C-D)**.

**Figure 4.**
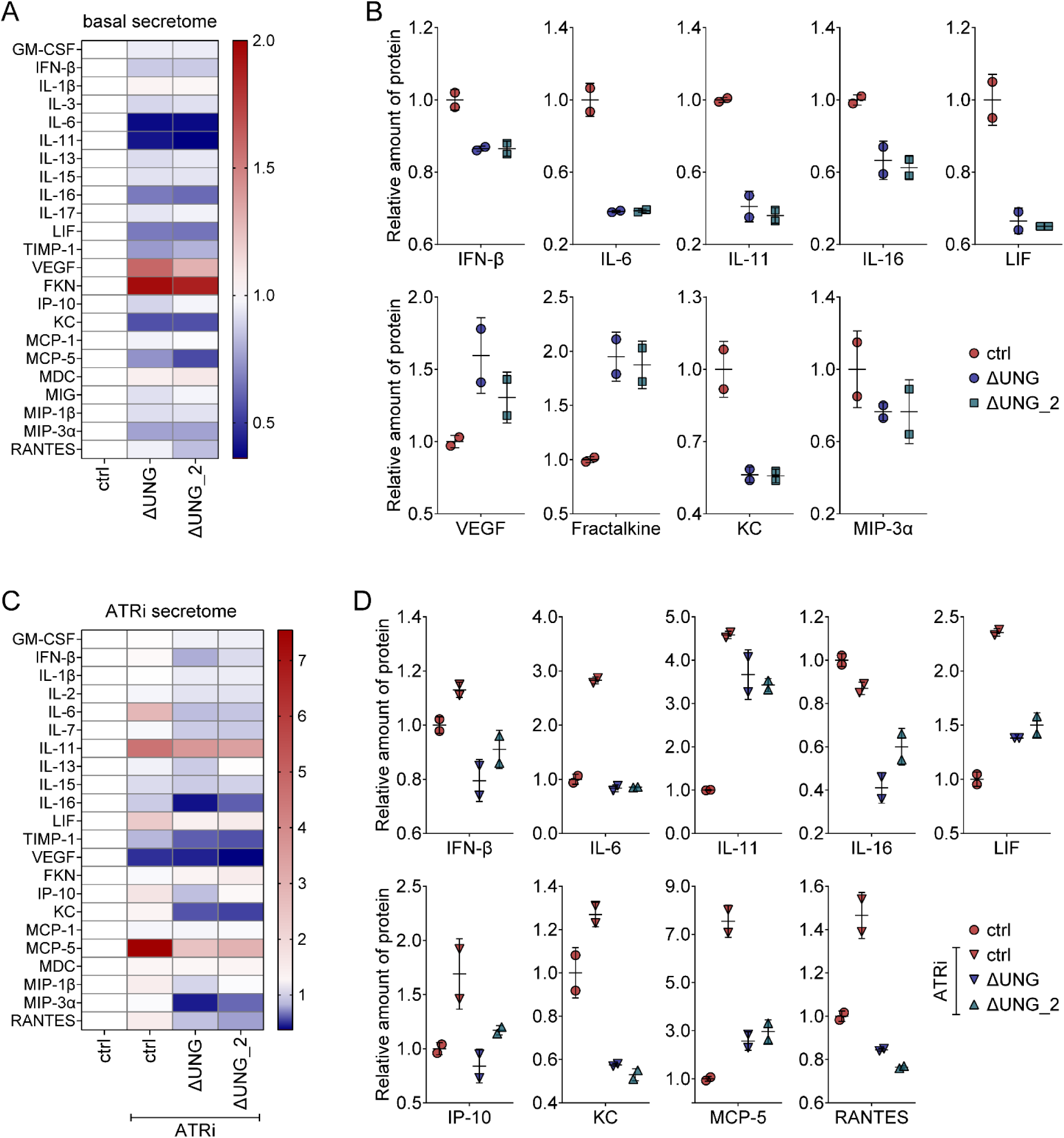
UNG deficiency alters the tumor cell secretome *in vitro*. **A**-**B.** Secretome analyses of vehicle-treated control (ctrl), ΔUNG, and ΔUNG_2 B16 tumor cells at 48 h. **A.** Heatmap showing mean relative amounts of a subset of secreted cytokines and chemokines (normalized to ctrl). **B.** Quantitation of the relative amounts of secreted IFN-β, IL-6, IL-11, IL-16, LIF, VEGF, Fractalkine, KC, and MIP-3α (normalized to mean of ctrl) at 48 h. Data from 1 experiment with 2 biological replicates. **C**-**D.** Secretome analyses of ctrl, ΔUNG, and ΔUNG_2 B16 tumor cells treated with 5 µM AZD6738 (ATRi) for 48 h, compared to vehicle-treated ctrl. **C.** Heatmap showing mean relative amounts of a subset of secreted cytokines and chemokines (normalized to ctrl) following ATRi treatment. **B.** Quantitation of the relative amounts of secreted IFN-β, IL-6, IL-11, IL-16, LIF, IP-10, KC, MCP-5, and RANTES (normalized to mean of ctrl) following ATRi treatment. Data from 1 experiment with 2 biological replicates. **B,D.** Means ± SD bars shown. Since n=2, no statistical tests were performed.

### UNG deficiency alters the cytokine and chemokine milieu *in vivo*

Our RNA-seq and secretome data show that UNG deficiency broadly represses inflammatory signaling pathways and reduces secretion of inflammatory cytokines and chemokines in B16 cells *in vitro*. We hypothesized that the suppression of tumor cell-intrinsic inflammatory signals would impact communication between tumor and immune cells and result in an altered cytokine and chemokine milieu in the tumor microenvironment (TME) *in vivo*. Using multiplex immunoassays, we examined inflammatory cytokines (GM-CSF, IFN-β, IFN-γ, IL-6, IL-12p70, IL-27p28, and TNF-α), and chemokines (IP-10, MCP-1, MCP-5, MIP-3α, and

RANTES) in TME. We first assayed these 12 analytes in extracts from control and ΔUNG B16 tumors harvested at days 6, 10, 14, and 18 post-tumor cell injection on day 1. In general, tumor cell-intrinsic UNG deficiency resulted in only modest changes in cytokines and chemokines in the TME, with minimal differences at days 6 and 18 **(Fig 5A)**. At day 10, on average, IFN-β and IP-10 were reduced while IFN-γ and TNF-α were elevated in ΔUNG tumors, although the differences did not reach statistical significance **(Fig 5A-B)**. At day 14, IFN-γ, TNF-α, and IP-10 were generally similar in control and ΔUNG tumors, but IL-6 and MIP-3α were significantly increased in ΔUNG tumors **(Fig 5C)**.

**Figure 5.**
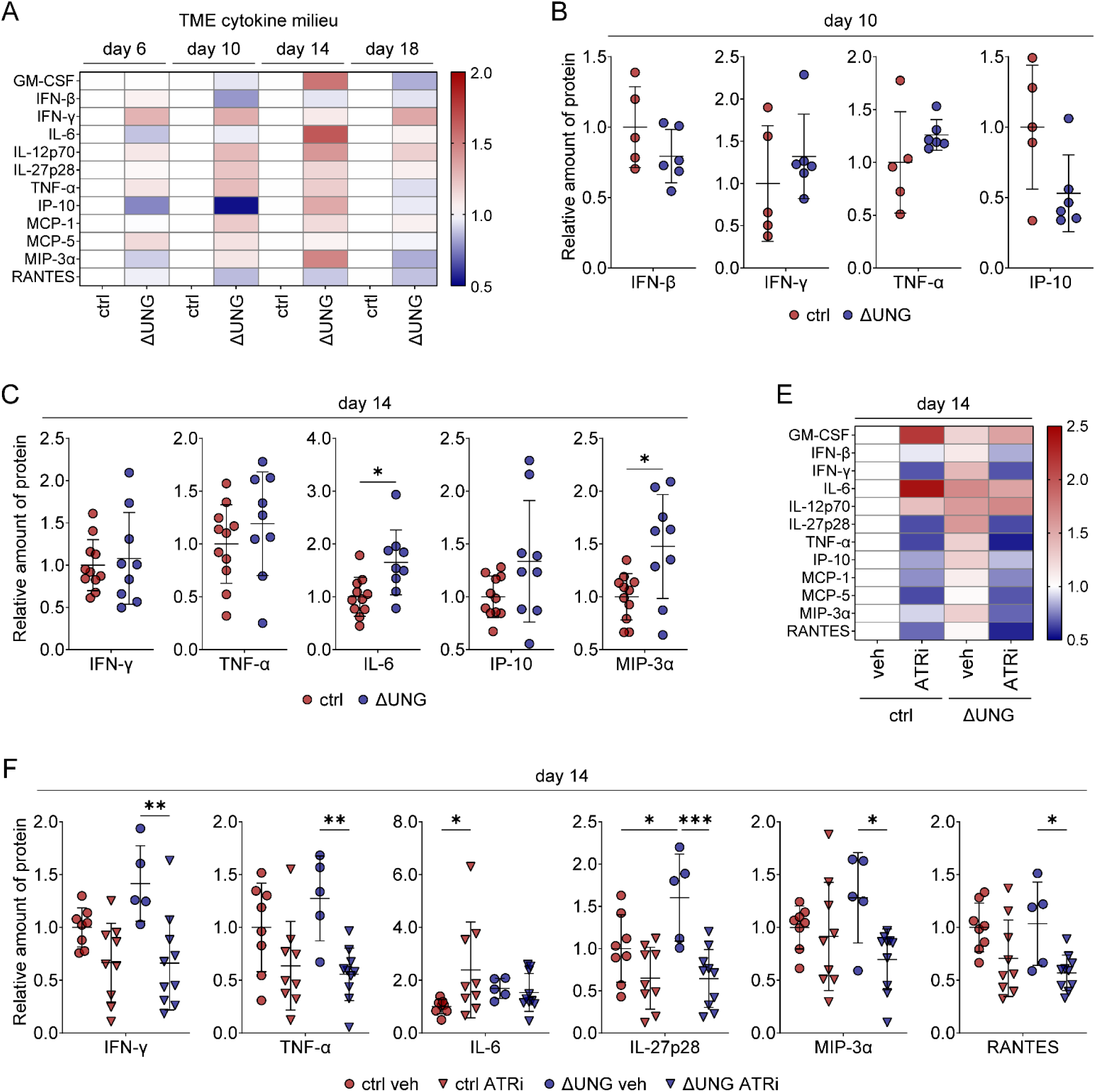
UNG deficiency alters the cytokine and chemokine milieu *in vivo*. **A**-**E.** Protein levels of 12 inflammatory cytokines and chemokines were measured in control (ctrl) and ΔUNG B16 tumors harvested from mice at the indicated time points. **A.** Heatmap depicting the mean relative amount of protein (normalized to ctrl for a given time point) in ctrl and ΔUNG B16 tumors harvested on days 6, 10, 14, and 18. **B.** Quantitation of the relative amounts of IFN-β, IFN-γ, TNF-α, and IP-10 (normalized to mean of ctrl) in tumors on day 10. Data from 1 experiment with n = 5 ctrl and 6 ΔUNG mice. No significant differences by two-tailed, unpaired Welch’s t test. **C.** Quantitation of the relative amounts of IFN-γ, TNF-α, IL-6, IP-10, and MIP-3α (normalized to mean of ctrl) in tumors on day 14. Data combined from 4 experiments with 1-4 mice per group each for total n = 11 ctrl and 9 ΔUNG. *p<0.05 by two-tailed, unpaired Welch’s t test. **D.** Heatmap depicting the mean relative amount of protein (normalized to ctrl veh) in a subset of ctrl and ΔUNG B16 tumors harvested on day 14 from mice treated with ATRi (75 mg/kg AZD6738, days 10-14) or vehicle (veh). **E.** Quantitation of the relative amounts of IFN-γ, TNF-α, IL-6, IL-27p28, MIP-3α and RANTES (normalized to mean of ctrl veh) at day 14. Data combined from 2 experiments with 2-6 mice per group each for total n = 8 ctrl veh, 5 ΔUNG veh, 9 ctrl ATRi, and 10 ΔUNG ATRi. *p<0.05, **p<0.01, ***p<0.001 by one-way ANOVA with Sidak’s multiple comparisons test. **B,C,E.** Mean ± SD bars shown.

Overall, levels of the assayed cytokine and chemokines in the TME were only modestly impacted by UNG deficiency in B16 tumor cells. We questioned whether treatment with ATRi would augment these differences, since our *in vitro* RNA-seq and secretome data show that ATRi-induced inflammatory signaling is attenuated in ΔUNG B16 cells. Therefore, we assayed the same 12 cytokines/chemokines in a subset of control and ΔUNG tumors harvested at day 14 from mice treated with vehicle or ATRi AZD6738 (75 mg/kg) once daily on days 10-14 **(Fig 5E)**. We chose day 14 for harvest as it was the only time point when we observed significant differences in untreated tumors. While ATRi treatment significantly induced IL-6 in control tumors, induction of IL-6 was absent in ΔUNG tumors **(Fig 5F),** just as induced secretion of IL-6 was absent from ATRi-treated ΔUNG cells *in vitro* (previously shown in Fig 4D). More striking was that ATRi-treated ΔUNG tumors had significantly decreased IFN-γ, TNF-α, IL-27p28, MIP-3α, and RANTES, compared to vehicle-treated ΔUNG tumors **(Fig 5F)**. The decreases in IFN-γ, TNF-α, and IL-27p28 in ΔUNG tumors were mirrored by average decreases of these cytokines in control tumors and are likely the result of direct effects of ATRi on immune cells. Conversely, decreases in MIP-3α and RANTES in ΔUNG tumors are likely, at least in part, tumor-cell specific, as we observed similar decreases in secretion of these chemokines *in vitro* following ATRi treatment (previously shown in Fig 4D).

### UNG deficiency alters tumor cell sensitivity to interferons *in vitro*

Our data demonstrate that loss of UNG in B16 tumor cells represses tumor-intrinsic inflammatory signaling pathways and the secretion of inflammatory cytokines and chemokines *in vitro*. UNG-deficient B16 tumors exhibit altered growth kinetics *in vivo* that are associated with changes in immune cell infiltration, notably reduced CD8^+^ T cells, greater CD4^+^ Tconv, and a more robust CD4^+^ Th1 response. Since differences in the quantities of assayed cytokines/chemokines in ΔUNG versus control tumors were modest, we hypothesized that altered sensitivity of ΔUNG tumor cells to cytokines may contribute to the observed differences in growth and the TME of ΔUNG tumors. We investigated whether ΔUNG B16 cells respond differently than control B16 cells to exogenous interferons *in vitro*. We treated cells for 18 hours with IFN-α (1.0 or 5.0 ng/mL) or IFN-β (0.2 or 1.0 ng/mL) and used upregulation of cell surface MHC-I and PD-L1 as readouts for responsiveness to these type I interferons **(Fig 6A-C; Supp Fig S7A)**. Control cells upregulated MHC-I significantly more in response to both IFN-α and IFN-β than did ΔUNG cells **(Fig 6B; Supp Fig S7A)** or ΔUNG_2 cells (**Supp Fig S7B)**. Upregulation of PD-L1 on control, ΔUNG, and ΔUNG_2 cells was similar following IFN-α treatment but was greater on ΔUNG and ΔUNG_2 cells treated 1.0 ng/mL IFN-β compared **(Fig 6C, Fig S7C)**.

**Figure 6.**
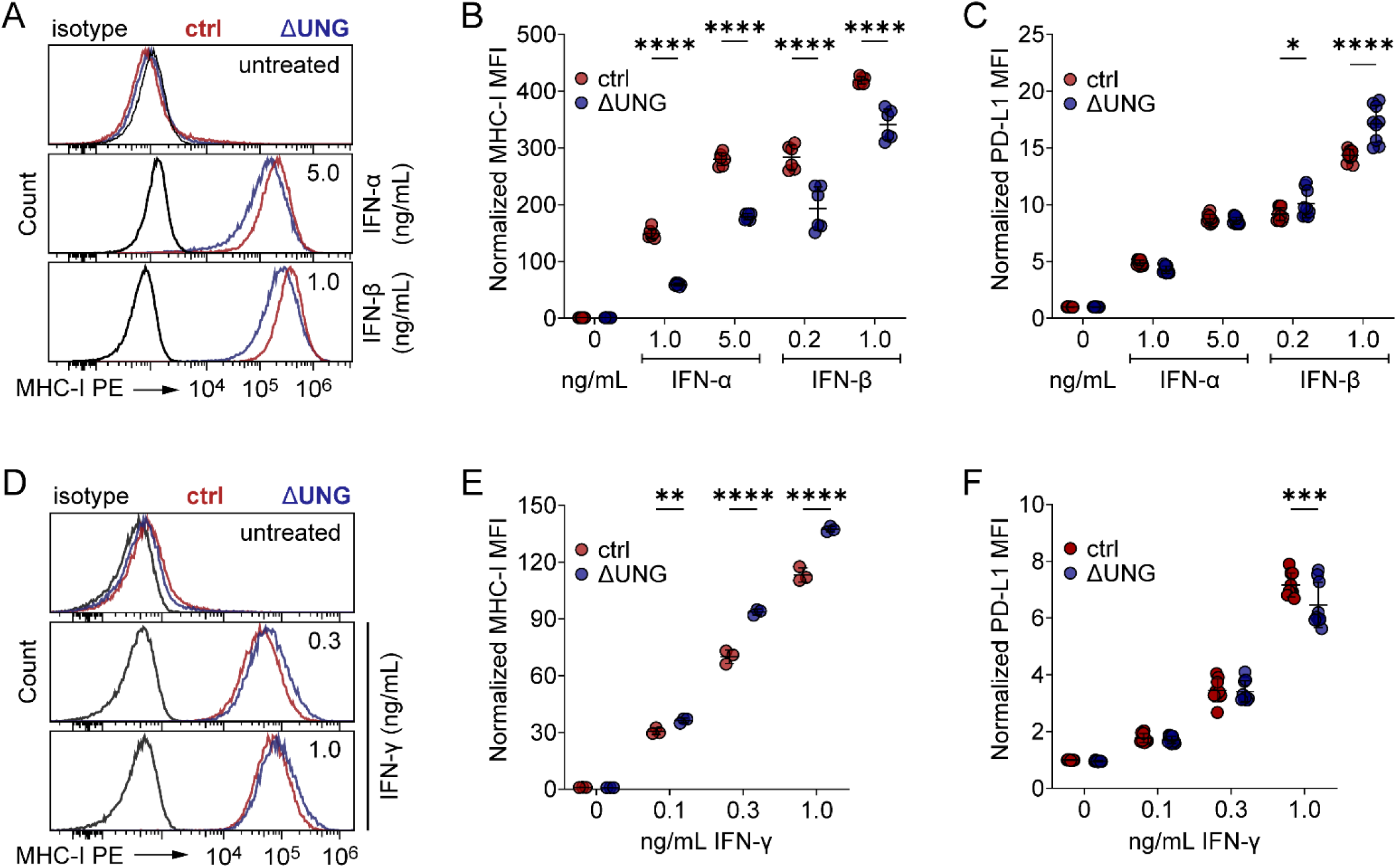
UNG deficiency alters tumor cell sensitivity to exogenous interferons *in vitro*. **A-C**. MHC-I and PD-L1 cell surface expression were analyzed on control (ctrl) and ΔUNG B16 cells treated *in vitro* with IFN-α (1.0 or 5.0 ng/mL) or IFN-β (0.2 or 1.0 ng/mL) for 18 h. **A.** Histograms showing the median fluorescence intensity (MFI) of MHC-I PE staining on representative untreated, IFN-α-treated (5.0 ng/mL), or IFN-β-treated (1.0 ng/mL) ctrl and ΔUNG B16 cells, compared to isotype control staining. **B-C.** Quantitation of MHC-I **(B)** and PD-L1 **(C)** MFI, normalized to the mean of untreated ctrl cells. Data from 2 representative (MHC-I) or 3 (PD-L1) independent experiments, each with 3 biological replicates, for n=6 (MHC-I) or n=9 (PD-L1) in total. Mean ± SD bars shown. *p<0.05, ****p<0.0001 by two-way ANOVA with Sidak’s multiple comparisons test. **D-F**. MHC-I and PD-L1 cell surface expression were analyzed on ctrl and ΔUNG B16 cells treated *in vitro* with IFN-γ (0.1, 0.3, or 1.0 ng/mL) for 18 h. **D.** Histograms showing the median fluorescence intensity (MFI) of MHC-I PE staining on representative untreated and IFN-γ-treated (0.3 or 1.0 ng/mL) ctrl and ΔUNG B16 cells, compared to isotype control staining. **E-F.** Quantitation MHC-I **(E)** and PD-L1 **(F)** MFI, normalized to the mean of untreated ctrl cells. Data from 1 representative (MHC-I) or 3 (PD-L1) independent experiments, each with 3 biological replicates, for n=3 (MHC-I) or n=9 (PD-L1) in total. Mean ± SD bars shown. **p<0.01,***p<0.001, ****p<0.0001 by two-way ANOVA with Tukey’s multiple comparisons test.

Next, we examined whether ΔUNG cells respond differently than control B16 cells to the type II interferon, IFN-γ (0.1, 0.3, or 1.0 ng/mL), again using upregulation of cell surface MHC-I and PD-L1 as readouts for responsiveness to IFN-γ **(Fig 6D-F; Supp Fig S7D)**. In contract to IFN-α/β, IFN-γ upregulated MHC-I significantly more on ΔUNG than control B16 cells **(Fig 6E; Supp Fig S7D)**. Upregulation of PD-L1 on control and ΔUNG cells was similar at the two lower doses of IFN-γ but was increased on control cells compared ΔUNG cells following treatment with 1.0 ng/mL IFN-γ **(Fig 6F)**. Regardless of the interferon used for treatment, the dynamic range in MHC-I upregulation between control and ΔUNG was much greater than for PD-L1 upregulation and may serve as the better readout for sensitivity to interferon in our models. Taken together, these data support that ΔUNG B16 cells are less sensitive to type I interferon but more sensitive to type II interferon than are control B16 cells.

## DISCUSSION

Here we show that murine B16 tumors deficient in uracil-DNA glycosylase, an enzyme that repairs dU lesions in DNA, have altered growth kinetics and immune cell infiltration in immunocompetent mice. In addition, UNG deficiency enhanced sensitivity of this checkpoint-resistant tumor model to anti-PD-L1 therapy. The *in vivo* differences we observed stem from a combination of repressed tumor cell-intrinsic signaling pathways, reduced secretion of inflammatory cytokines and chemokines from tumor cells and the resultant changes in communication with host immune cells, as well as altered sensitivity of ΔUNG tumor cells to exogenous type-I and type-II interferons.

B16 tumor cell-specific UNG deficiency significantly impacted basal and ATR-induced inflammatory signaling *in vitro*. This was evident in both the transcriptome and the secretome. Yet the impacts of tumor cell-specific UNG deficiency *in vivo* were relatively modest. However, the observed delays in early tumor growth were highly reproducible and were clearly immune system-dependent, as no differences in growth were seen in tumors in athymic nude mice that lack mature T cells. Consistent with this, we observed altered T cell infiltration on day 14 in UNG-deficient tumors, with a shift toward reduced CD8^+^ T cell and increased CD4^+^ T cell infiltration, independent of changes in regulatory T cells (Treg). Instead, UNG-deficient tumors had increased conventional CD4^+^ T cells (CD4^+^ Tconv), including CD4^+^ Th1 effector cells. Th1 cells are inflammatory CD4^+^ effector T cells that secrete IFN-γ (and often TNF-α) to activate cytolytic activity of macrophages and CD8^+^ cytotoxic T lymphocytes (37–40). Th1 cells also facilitate direct tumor cell killing via cytokine release and activation of death receptors on tumor cells (37, 40). Th1 responses are associated with improved response to immune checkpoint blockade in mice (41), as well as better outcomes in patients across multiple cancer types (42). The Th1 response is driven by differentiation of CD4^+^ Tconv following presentation of MHC-II bound antigen by antigen-presenting cells (APC) in the presence of IFN-γ, IL-12, and IL-27 (43–45), and we observed increased intratumoral IL-27p28 on day 14 in vehicle-treated UNG-deficient tumors. Taken together, our data support that UNG-deficient tumors promote a stronger Th1 response than control tumors. However, the CD8^+^ T cell response was blunted in UNG-deficient tumors, and while the relative CD8^+^ T cell cytokine competency was unchanged, UNG-deficient tumors have reduced CD8^+^ T cell infiltration overall. Consistent with this, UNG-deficient tumors had reduced CD103^+^ dendritic cells (DC) on day 14, which are important APC for priming the CD8^+^ T cell response (28). The interplay of these negative and positive changes in DC and T cell infiltration likely underlies why UNG-deficient tumors exhibited early growth delay that was not durable, with growth accelerating rapidly after day 17.

Analyses of inflammatory cytokines and chemokines in tumors at multiple time points yielded only subtle differences in IFN-β, IFN-γ, and other cytokines and chemokines. This is likely because many cytokines and chemokines in the TME originate from non-tumor cell sources, including infiltrating immune cells and tumor-associated, non-immune cells (46–48). However, large changes in some of these cytokines and chemokines may not be necessary to alter the microenvironment in UNG-deficient tumors. UNG deficiency clearly altered interferon-dependent upregulation of cell surface MHC-I *in vitro*, with UNG-deficient cells being less responsive to type-I interferon (IFN-α/β) and more responsive to type-II interferon (IFN-γ). Interferons from early infiltrating innate immune cells may differentially alter MHC-I expression in UNG-deficient versus control cells, which may then impact anti-tumor activity of NK and CD8^+^ T cells (49–55). Further investigation of how UNG-deficient, as well as UNG-inhibited, cells respond to different cytokine and chemokine signals is warranted.

Our study utilized a single model system (B16 melanoma) to limit confounding factors outside of our primary variable, UNG activity. The study is limited in that only the tumor cells were deficient in UNG activity, whereas UNG inhibitors may have direct and important consequences in immune cells. In addition, UNG inhibitors may not phenocopy *Ung* knockout systems. Bioavailable UNG inhibitors are necessary to address these limitations using *in vivo* models. Overall, our data provide proof-of-concept evidence that UNG inhibitors may modulate anti-tumor immune responses and sensitize tumors to immune checkpoint blockade in certain cancer contexts. This may be of particular relevance for therapeutic regimens involving standard-of-care antimetabolites, including 5-fluorouracil, pemetrexed, cytarabine, hydroxyurea, fludarabine, gemcitabine, methotrexate, 6-mercaptopurine, and capecitabine, that disrupt nucleotide pools and increase dU incorporation by DNA polymerases. Patients with low tumor UNG levels may benefit more from antimetabolites combined with checkpoint inhibitors, while patients with high tumor UNG levels may be candidates for UNG inhibitor therapy.

## METHODS

### Cell lines

B16-F10 (ATCC, referred to as B16 throughout) were cultured in DMEM, 10% fetal bovine serum, 1% Pen/Strep in a humidified cell culture incubator set to 37°C, 5% CO_2_. Two clonal populations of B16 *Ung* knockout cells (ΔUNG and ΔUNG_2) and a clonal population of Cas9-expressing control B16 cells (referred to as control in text, ctrl in figures) were generated and validated in this study (**Supp Fig S1 and S2**). Cell lines were periodically tested for mycoplasma contamination using the MycoStrip Mycoplasma Detection Kit (InvivoGen). Cells under passage number 15 post-selection were used for all experiments. Control, ΔUNG, and ΔUNG_2 cells were maintained under 1.5 µg/mL puromycin (selection condition) during routine passaging. Cell line treatments were performed in complete media without puromycin.

### Animal experiments

Experiments were performed in accordance with protocols approved by the University of Pittsburgh Animal Care and Use Committee. Control and ΔUNG B16 cells (2.5 x 10^5^) in serum-free DMEM were subcutaneously injected into the right hind flank of 8–10-week-old C57BL/6 mice or athymic nude mice (purchased from Jackson Laboratories). Tumors were measured with digital calipers on the indicated days, and volumes calculated as volume = (length x width2)/2. For studies comparing the rates of growth of control and ΔUNG B16 in C57BL/6 mice or nude mice, only tumors that established were included in measurements. For the study involving anti-PD-L1 therapy, mice were injected intraperitoneally with 100 µg anti-PD-L1 antibody (clone 10F.9G2, BioXCell inVivoPlus) diluted in 100 µL inVivoPure pH 6.5 Dilution Buffer (BioXCell), every 3 days for 6 doses, starting on day 2 after tumor cell injection on day 1. Vehicle-treated mice were injected intraperitoneally with 100 µL inVivoPure pH 6.5 Dilution Buffer. Since treatment began the day after tumor cell injection, all tumors, including those that did not take or that completely resolved, were included in the study group measurements. The tumor endpoint was reached when tumor volume exceeded 1000 mm^3^ or the tumor ulcerated. When multiple tumors within a group reached their endpoint, the entire study group was deemed to have reached the experimental endpoint. Mice that were euthanized due to tumor ulceration prior to the experimental endpoint were excluded from the reported tumor volumes.

### Flow cytometry for immune profiling

Control or ΔUNG B16 tumors and spleens were harvested at the indicated time points. Splenocytes were used for single color controls, fluorescence-minus-one (FMO) controls, and for general gating. Tumor tissue (≤ 250 mg) was minced and then digested in Collagenase IV Cocktail containing Collagenase IV, DNase I, Soybean Trypsin Inhibitor (all Worthington). Tumor homogenate was smashed through a 70 μm cell strainer using the rubber plunger of a syringe. Spleens were mechanically dissociated between frosted glass slides and filtered through 70 μm cell strainers. Erythrocytes were lysed and cell suspensions were counted with a Scepter 3.0 (Millipore) and seeded at 1.2-2 x 10^6^ cells in 96-well round bottom plates for blocking and staining as follows: Fc receptors were blocked with anti-CD16/32 antibody, cells were stained with antibodies to surface antigens and eFlour780 viability dye or LIVE/DEAD fixable near-IR dead cell stain, samples were fixed and permeabilized in eBioscience Fixation/Permeabilization reagent (Invitrogen), and when performing nuclear (Ki67, Foxp3) or intracellular cytokine (IFN-γ, TNF-α, IL-17) staining, samples were stained with antibodies to nuclear/intracellular proteins. Antibody information (target, conjugate, clone, dilution) can be found in the key resources table in the supplemental materials. Brilliant Stain Buffer Plus (BD Biosciences) was added to antibody cocktails containing multiple Brilliant Violet dye conjugates to prevent polymer dye-dye interactions. For measurement of cytokine-producing CD8^+^ and CD4^+^ T cells, prior to staining, cells were stimulated for 4 h with 1x eBioscience Cell Stimulation Cocktail (PMA, ionomycin, and protein transport inhibitors; Invitrogen) in complete DMEM media. Unstimulated controls were treated with 1x eBioscience Cell Protein Transport Inhibitors cocktail (Invitrogen). Since PMA/ionomycin stimulation induces internalization of CD3 and the CD8/CD4 co-receptors, staining for CD3, CD4, CD8, and CD45 was performed post-fixation/permeabilization during intracellular cytokine staining.

Data were acquired uncompensated data using a BD LSRFortessa 4-laser cytometer and BD FACSDiva software, with compensation performed in FlowJo V10 software, or data were acquired compensated/unmixed using a Cytek Aurora spectral cytometer and SpectroFlow software. Single stained spleen samples with matching unstained cells or single stained OneComp eBeads or UltraComp eBeads Plus (Invitrogen) were used for single color compensation controls. Fluorescence-minus-one (FMO) controls were used, where appropriate, to empirically determine gating. All data analyses were performed in FlowJo V10 software. Gating strategies are shown in Supplemental Figures S8-S12.

### RNA-seq and data analysis

Control, ΔUNG, and ΔUNG_2 B16 cells were treated with 5 µM ATRi AZD6738 (provided by AstraZeneca) or 0.05% DMSO vehicle for 48 h. Cells were harvested in Trizol and total RNA was extracted using the Direct Zol RNA kit (Zymo Research Corp). RNA was submitted to the Novogen Advancing Genomics facility (Sacramento, CA, USA) for RNA sequence analysis. Read alignment was performed using HISAT2 software using the mouse reference genome. FPKM was used to quantify the abundance of transcripts or genes. Differential gene expression analysis was performed using DEseq2 (for biological replicates) or edgeR (for no biological replicates) software with FDR correction by the Benjamini-Hochberg procedure with threshold [log2(foldchange)]>=1 & p-adj <=0.05. Read alignment and differential gene expression analysis were performed by Novogen. Gene set enrichment analysis (GSEA 4.3.2) was utilized to perform rank-based identification of the most enriched pathways between groups. Group pathway analysis was performed using ClusterProfiler software for Gene Ontology analysis (http://www.geneontology.org/). GO terms with p-adj <0.05 are significantly enriched, and the most significant terms were selected for display.

### *In vitro* secretome analyses

Control, ΔUNG, and ΔUNG_2 B16 cell were treated in vitro with vehicle or 5 µM AZD6738 (ATRi) for 48 h. Media from treated cells was collected, centrifuged (350 x g), and supernatant was frozen at −80°C. Frozen supernatants were sent on dry ice to Eve Technologies for analyses using their Mouse Cytokine/Chemokine 44-Plex Discovery Assay Array.

### Measurement of intratumoral cytokines and chemokines

Control and ΔUNG B16 cells (1.0 x 10^5^ for day 6 harvest, 5 x 10^5^ for day 10 harvest, or 2.5 x 10^5^ for day 14 or 18 harvest) in serum-free DMEM were subcutaneously injected into the right hind flank of 8–10-week-old C57BL/6 mice or athymic nude mice (purchased from Jackson Laboratories). Control and ΔUNG tumors were harvested at the indicated time points, portioned, and frozen on dry ice. For a subset of day 14 samples, mice were treated with ATRi AZD6738 (75 mg/kg) or vehicle (10% DMSO, 40% Propylene Glycol, 50% dH_2_O) once daily on days 10-14, and tumors were harvested 2 h after the last dose on day 14. Frozen tumor pieces were weighed, and >20 mg (but <70mg) of tumor was added to Precellys CK28-R Protein Safe Hard tissue homogenizing tubes (Bertin Technologies) containing Invitrogen ProcartaPlex Cell Lysis Buffer (Thermo Fisher Scientific) with 1 mM PMSF at a volume of 500 μL per 75 mg tumor. To generate protein extracts, samples were homogenized using a Precellys 24 homogenizer (Bertin Instruments) (2 cycles of 6,000 rpm × 15 seconds, samples on ice ≥2 minutes, 2 additional cycles of 6,000 rpm × 15 seconds) and were cleared of insoluble material. Aliquots were stored at −80°C. Protein levels of 12 analytes (GM-CSF, IFN-β, IFN-γ, IL-6, IL-12p70, IL-27p28, TNF-α, IP-10, MCP-1, MCP-5, MIP-3α, and RANTES) in tumor extracts were determined using the U-Plex platform and MESO QuickPlex SQ 120 (Mesoscale Discovery) according to the manufacturer’s instructions, with the addition of overnight incubation (4°C) of samples with the capture antibodies. Tumor extracts were diluted 1:10 in assay diluent for MCP-1 and RANTES, 1:50 in assay diluent for IP-10 and MCP-5, or were assayed undiluted for the remaining analytes. Data analyses were performed in MSD Discovery Workbench software (Mesoscale Discovery), and protein concentrations (pg/mL) were normalized to the mean concentrations of vehicle control samples for a given target at a given time point, assayed on the same plate, and the data are presented as the relative amount of protein compared with vehicle control. No inter-plate or inter-run comparisons were made.

### Flow cytometry for MHC-I and PD-L1 expression *in vitro*

Cells were treated with 1.0 or 5.0 ng/mL recombinant mouse IFN-α1 (BioLegend), 0.2 or 1.0 ng/mL recombinant mouse IFN-β1 (BioLegend), 0.1, 0.3, or 1.0 ng/mL recombinant mouse IFN-γ (Peprotech, dissolved according to the manufacturer’s instructions), or equivalent volumes of 1x PBS, for 18 h. Interferon-treated cells were harvested using Accutase (Gibco) cell dissociation reagent (IFN-α/β experiments) or were gently scraped for collection (IFN-γ experiments). Cells were blocked in 5% normal mouse serum, stained with MHC-I or PD-L1 or isotype control antibodies, and efluor780 fixable viability dye. Cells were analyzed live or were fixed in 175 µL FluoroFix (BioLegend) prior to analyses. Single-stained OneComp eBeads (Invitrogen) were used as compensation controls for MHC-I and PD-L1. A single-stained mix of live and heat-killed (30 sec at 95°C) cells were used as the compensation control for efluor780. Acquisition was performed with a 4-laser CytoFLEX (Beckman Coulter), and analyses were performed in FlowJo V10. The gating strategy is shown in Supplemental Figure S13.

### Statistics

The number of independent experiments and biological replicates (or mice), and the statistical tests performed (in GraphPad Prism 10) are specified in figure legends. A 95% confidence interval and significance of P less than 0.05 were used for all statistical tests. Data are reported as mean ± SD except for tumor volume data, which is presented as mean ± SEM. Stat bars are shown only for comparisons that were statistically significant (unless otherwise specified), but significance is adjusted for all comparisons made when multiple comparisons tests were performed.

## ACKNOWLEDGEMENTS

This work was supported by CA236367 and CA266172 (CJB), CA148629, ES029518, ES028949, CA238061, and ES032522 (RWS) from the NIH. This project used the Cytometry Facility that is supported in part by award P30CA047904 from the NIH. Support was also provided by the Legoretta Cancer Center Endowment Fund (RWS). We thank Chris Koczor and Jianfeng Li (University of South Alabama) for discussions at the start of this project.

## AUTHOR CONTRIBUTIONS

FPV, PP, CRS, and REJ designed and completed experiments in the Bakkenist lab. FPV, CRS, and PP completed the *in vivo* experiments. DI generated lentivirus and performed the molecular beacon assay for uracil DNA glycosylase activity in the Sobol lab. DI and RWS analyzed the molecular beacon assay data (Supplemental Figure S2). All authors contributed to writing the paper.

## REFERENCES

1. Lindahl T. Instability and decay of the primary structure of DNA. Nature. 1993;362(6422):709–15.

2. Alexandrov LB, Nik-Zainal S, Wedge DC, Aparicio SA, Behjati S, Biankin AV, et al. Signatures of mutational processes in human cancer. Nature. 2013;500(7463):415–21.

3. Roberts SA, Lawrence MS, Klimczak LJ, Grimm SA, Fargo D, Stojanov P, et al. An APOBEC cytidine deaminase mutagenesis pattern is widespread in human cancers. Nat Genet. 2013;45(9):970–6.

4. Isozaki H, Sakhtemani R, Abbasi A, Nikpour N, Stanzione M, Oh S, et al. Therapy-induced APOBEC3A drives evolution of persistent cancer cells. Nature. 2023;620(7973):393–401.

5. Rose MG, Farrell MP, and Schmitz JC. Thymidylate synthase: a critical target for cancer chemotherapy. Clin Colorectal Cancer. 2002;1(4):220–9.

6. Sarno A, Lundbaek M, Liabakk NB, Aas PA, Mjelle R, Hagen L, et al. Uracil-DNA glycosylase UNG1 isoform variant supports class switch recombination and repairs nuclear genomic uracil. Nucleic Acids Res. 2019;47(9):4569–85.

7. Nilsen H, Rosewell I, Robins P, Skjelbred CF, Andersen S, Slupphaug G, et al. Uracil-DNA glycosylase (UNG)-deficient mice reveal a primary role of the enzyme during DNA replication. Mol Cell. 2000;5(6):1059–65.

8. Kavli B, Iveland TS, Buchinger E, Hagen L, Liabakk NB, Aas PA, et al. RPA2 winged-helix domain facilitates UNG-mediated removal of uracil from ssDNA; implications for repair of mutagenic uracil at the replication fork. Nucleic Acids Res. 2021;49(7):3948–66.

9. Almeida KH, and Sobol RW. A unified view of base excision repair: lesion-dependent protein complexes regulated by post-translational modification. DNA Repair (Amst). 2007;6(6):695–711.

10. Saxena S, Nabel CS, Seay TW, Patel PS, Kawale AS, Crosby CR, et al. Unprocessed genomic uracil as a source of DNA replication stress in cancer cells. Mol Cell. 2024;84(11):2036–52 e7.

11. Mortusewicz O, Haslam J, Gad H, and Helleday T. Uracil-induced replication stress drives mutations, genome instability, anti-cancer treatment efficacy, and resistance. Mol Cell. 2025;85(10):1897–906.

12. Musiani D, Yucel H, Vallette M, Angrisani A, El Botty R, Ouine B, et al. Uracil processing by SMUG1 in the absence of UNG triggers homologous recombination and selectively kills BRCA1/2-deficient tumors. Mol Cell. 2025;85(6):1072–84 e10.

13. Sugitani N, Vendetti FP, Cipriano AJ, Pandya P, Deppas JJ, Moiseeva TN, et al. Thymidine rescues ATR kinase inhibitor-induced deoxyuridine contamination in genomic DNA, cell death, and interferon-α/β expression. Cell Reports. 2022;40(12):111371.

14. Savva R. Targeting uracil-DNA glycosylases for therapeutic outcomes using insights from virus evolution. Future Med Chem. 2019;11(11):1323–44.

15. Grundy GJ, and Parsons JL. Base excision repair and its implications to cancer therapy. Essays Biochem. 2020;64(5):831–43.

16. Mechetin GV, Endutkin AV, Diatlova EA, and Zharkov DO. Inhibitors of DNA Glycosylases as Prospective Drugs. Int J Mol Sci. 2020;21(9).

17. Malfatti MC, Bellina A, Antoniali G, and Tell G. Revisiting Two Decades of Research Focused on Targeting APE1 for Cancer Therapy: The Pros and Cons. Cells. 2023;12(14).

18. Zhang P, He F, and Chang X. Single G-quadruplex-based fluorescence method for the uracil-DNA glycosylase inhibitor screening. Heliyon. 2024;10(17):e37171.

19. Montaldo NP, Nilsen HL, and Bordin DL. Targeting base excision repair in precision oncology. DNA Repair (Amst). 2025;149:103844.

20. Bulgar AD, Weeks LD, Miao Y, Yang S, Xu Y, Guo C, et al. Removal of uracil by uracil DNA glycosylase limits pemetrexed cytotoxicity: overriding the limit with methoxyamine to inhibit base excision repair. Cell Death Dis. 2012;3(1):e252.

21. Weeks LD, Fu P, and Gerson SL. Uracil-DNA glycosylase expression determines human lung cancer cell sensitivity to pemetrexed. Mol Cancer Ther. 2013;12(10):2248–60.

22. Weeks LD, Zentner GE, Scacheri PC, and Gerson SL. Uracil DNA glycosylase (UNG) loss enhances DNA double strand break formation in human cancer cells exposed to pemetrexed. Cell Death Dis. 2014;5(2):e1045.

23. Lu CS, Lin CW, Chang YH, Chen HY, Chung WC, Lai WY, et al. Antimetabolite pemetrexed primes a favorable tumor microenvironment for immune checkpoint blockade therapy. J Immunother Cancer. 2020;8(2).

24. Principe N, Phung AL, Stevens KLP, Elaskalani O, Wylie B, Tilsed CM, et al. Anti-metabolite chemotherapy increases LAG-3 expressing tumor-infiltrating lymphocytes which can be targeted by combination immune checkpoint blockade. J Immunother Cancer. 2024;12(9).

25. Chen S, Lee LF, Fisher TS, Jessen B, Elliott M, Evering W, et al. Combination of 4-1BB agonist and PD-1 antagonist promotes antitumor effector/memory CD8 T cells in a poorly immunogenic tumor model. Cancer Immunol Res. 2015;3(2):149–60.

26. Kleffel S, Posch C, Barthel SR, Mueller H, Schlapbach C, Guenova E, et al. Melanoma Cell-Intrinsic PD-1 Receptor Functions Promote Tumor Growth. Cell. 2015;162(6):1242–56.

27. Lin H, Wei S, Hurt EM, Green MD, Zhao L, Vatan L, et al. Host expression of PD-L1 determines efficacy of PD-L1 pathway blockade-mediated tumor regression. J Clin Invest. 2018;128(2):805–15.

28. del Rio ML, Rodriguez-Barbosa JI, Kremmer E, and Forster R. CD103- and CD103+ bronchial lymph node dendritic cells are specialized in presenting and cross-presenting innocuous antigen to CD4+ and CD8+ T cells. J Immunol. 2007;178(11):6861–6.

29. Dovedi SJ, Adlard AL, Lipowska-Bhalla G, McKenna C, Jones S, Cheadle EJ, et al. Acquired resistance to fractionated radiotherapy can be overcome by concurrent PD-L1 blockade. Cancer Res. 2014;74(19):5458–68.

30. Abiko K, Matsumura N, Hamanishi J, Horikawa N, Murakami R, Yamaguchi K, et al. IFN-gamma from lymphocytes induces PD-L1 expression and promotes progression of ovarian cancer. Br J Cancer. 2015;112(9):1501–9.

31. Qian J, Wang C, Wang B, Yang J, Wang Y, Luo F, et al. The IFN-gamma/PD-L1 axis between T cells and tumor microenvironment: hints for glioma anti-PD-1/PD-L1 therapy. J Neuroinflammation. 2018;15(1):290.

32. Vendetti FP, Karukonda P, Clump DA, Teo T, Lalonde R, Nugent K, et al. ATR kinase inhibitor AZD6738 potentiates CD8+ T cell-dependent antitumor activity following radiation. J Clin Invest. 2018;128(9):3926–40.

33. Dillon MT, Bergerhoff KF, Pedersen M, Whittock H, Crespo-Rodriguez E, Patin EC, et al. ATR Inhibition Potentiates the Radiation-induced Inflammatory Tumor Microenvironment. Clin Cancer Res. 2019;25(11):3392–403.

34. Feng X, Tubbs A, Zhang C, Tang M, Sridharan S, Wang C, et al. ATR inhibition potentiates ionizing radiation-induced interferon response via cytosolic nucleic acid-sensing pathways. EMBO J. 2020;39(14):e104036.

35. Vendetti FP, Pandya P, Clump DA, Schamus-Haynes S, Tavakoli M, diMayorca M, et al. The schedule of ATR inhibitor AZD6738 can potentiate or abolish antitumor immune responses to radiotherapy. JCI Insight. 2023;8(4).

36. Hardaker EL, Sanseviero E, Karmokar A, Taylor D, Milo M, Michaloglou C, et al. The ATR inhibitor ceralasertib potentiates cancer checkpoint immunotherapy by regulating the tumor microenvironment. Nat Commun. 2024;15(1):1700.

37. Knutson KL, and Disis ML. Tumor antigen-specific T helper cells in cancer immunity and immunotherapy. Cancer Immunol Immunother. 2005;54(8):721–8.

38. Bos R, and Sherman LA. CD4+ T-cell help in the tumor milieu is required for recruitment and cytolytic function of CD8+ T lymphocytes. Cancer Res. 2010;70(21):8368–77.

39. Eisel D, Das K, Dickes E, Konig R, Osen W, and Eichmuller SB. Cognate Interaction With CD4(+) T Cells Instructs Tumor-Associated Macrophages to Acquire M1-Like Phenotype. Front Immunol. 2019;10:219.

40. Montauti E, Oh DY, and Fong L. CD4(+) T cells in antitumor immunity. Trends Cancer. 2024;10(10):969–85.

41. Lauret Marie Joseph E, Kirilovsky A, Lecoester B, El Sissy C, Boullerot L, Rangan L, et al. Chemoradiation triggers antitumor Th1 and tissue resident memory-polarized immune responses to improve immune checkpoint inhibitors therapy. J Immunother Cancer. 2021;9(7).

42. Fridman WH, Pages F, Sautes-Fridman C, and Galon J. The immune contexture in human tumours: impact on clinical outcome. Nat Rev Cancer. 2012;12(4):298–306.

43. Pflanz S, Timans JC, Cheung J, Rosales R, Kanzler H, Gilbert J, et al. IL-27, a heterodimeric cytokine composed of EBI3 and p28 protein, induces proliferation of naive CD4+ T cells. Immunity. 2002;16(6):779–90.

44. Zhu J, Yamane H, and Paul WE. Differentiation of effector CD4 T cell populations (*). Annu Rev Immunol. 2010;28:445–89.

45. Lee J, Lozano-Ruiz B, Yang FM, Fan DD, Shen L, and Gonzalez-Navajas JM. The Multifaceted Role of Th1, Th9, and Th17 Cells in Immune Checkpoint Inhibition Therapy. Front Immunol. 2021;12:625667.

46. Swiecki M, and Colonna M. Type I interferons: diversity of sources, production pathways and effects on immune responses. Curr Opin Virol. 2011;1(6):463–75.

47. Boukhaled GM, Harding S, and Brooks DG. Opposing Roles of Type I Interferons in Cancer Immunity. Annu Rev Pathol. 2021;16:167–98.

48. Yu R, Zhu B, and Chen D. Type I interferon-mediated tumor immunity and its role in immunotherapy. Cell Mol Life Sci. 2022;79(3):191.

49. Elliott JM, and Yokoyama WM. Unifying concepts of MHC-dependent natural killer cell education. Trends Immunol. 2011;32(8):364–72.

50. He Y, and Tian Z. NK cell education via nonclassical MHC and non-MHC ligands. Cell Mol Immunol. 2017;14(4):321–30.

51. Paul S, and Lal G. The Molecular Mechanism of Natural Killer Cells Function and Its Importance in Cancer Immunotherapy. Front Immunol. 2017;8:1124.

52. Bern MD, Parikh BA, Yang L, Beckman DL, Poursine-Laurent J, and Yokoyama WM. Inducible down-regulation of MHC class I results in natural killer cell tolerance. J Exp Med. 2019;216(1):99–116.

53. Taylor BC, and Balko JM. Mechanisms of MHC-I Downregulation and Role in Immunotherapy Response. Front Immunol. 2022;13:844866.

54. Lerner EC, Woroniecka KI, D’Anniballe VM, Wilkinson DS, Mohan AA, Lorrey SJ, et al. CD8(+) T cells maintain killing of MHC-I-negative tumor cells through the NKG2D-NKG2DL axis. Nat Cancer. 2023;4(9):1258–72.

55. Mengoni M, Mahlo FO, Gaffal E, Tuting T, and Braun AD. Downregulation of MHC-I on Melanoma Cells and Decreased CD8+ T-Cell Infiltration Are Associated With Metastatic Spread and Resistance to Immunotherapy. Lab Invest. 2025;105(3):102209.

