## SUPPLEMENTAL DATA for "Uracil-DNA glycosylase deficiency is associated with repressed tumor cell-intrinsic inflammatory signaling and altered sensitivity to exogenous interferons"

### Nucleotide sequence alignment

cas9-expressing control (wildtype UNG)

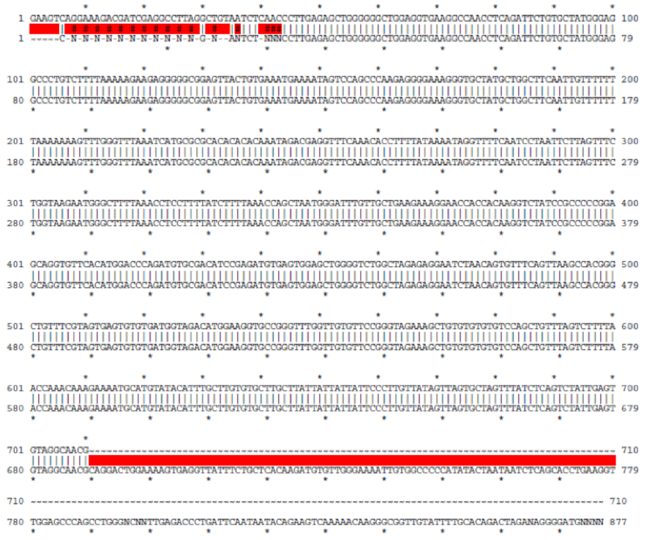

ΔUNG

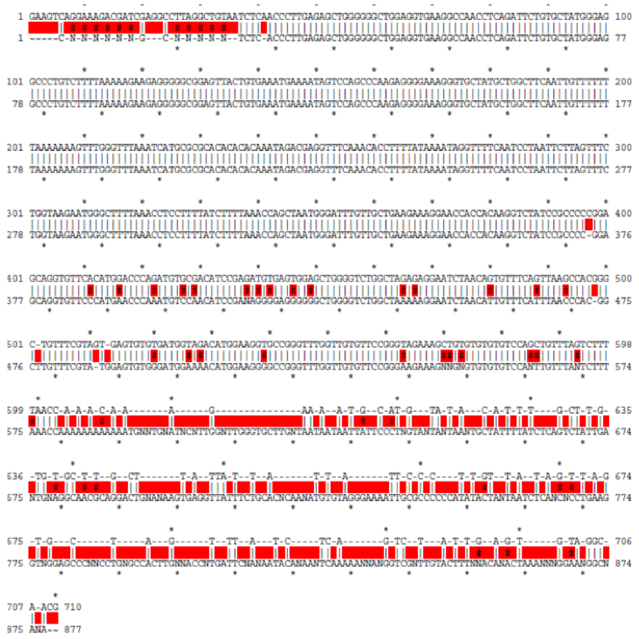

ΔUNG_2

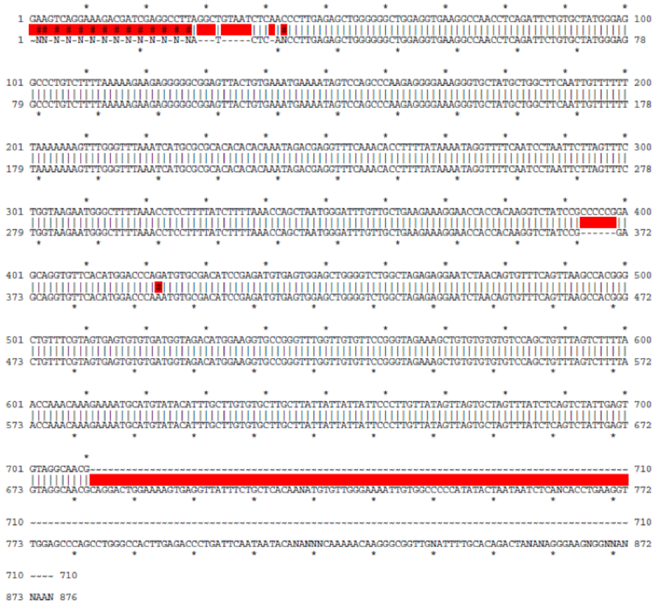

### Amino acid sequence alignment

cas9-expressing control

Leu Met Gly Phe Val Ala Glu Glu Arg Asn His His Lys Val Tyr Pro Pro Pro
Glu Gln Val Phe Thr Trp Thr Gln Met Cys Asp Ile Arg Asp

ΔUNG

*Changes in AA*

Leu Met Gly Phe Val Ala Glu Glu Arg Asn His His Lys Val Tyr Pro Pro Arg
Ser Arg Cys Ser His Glu Pro Lys Cys Pro Thr Ser Xxx Arg Gly Gly Gly

ΔUNG_2

*2 Proline deletion*

Leu Met Gly Phe Val Ala Glu Glu Arg Asn His His Lys Val Tyr Pro Glu Gln Val
Phe Thr Trp Thr Gln Met Cys Asp Ile Arg Asp

**Supplemental Figure S1. Sanger sequence and amino acid alignment for *Ung* knockout B16 cell lines.** Two clonal, homozygous *Ung* knockout B16 cell lines (named ΔUNG and ΔUNG_2) were used in this study. ΔUNG has a homozygous deletion that generates a frameshift and a stop codon. ΔUNG_2 has a homozygous deletion that deletes two proline residues.

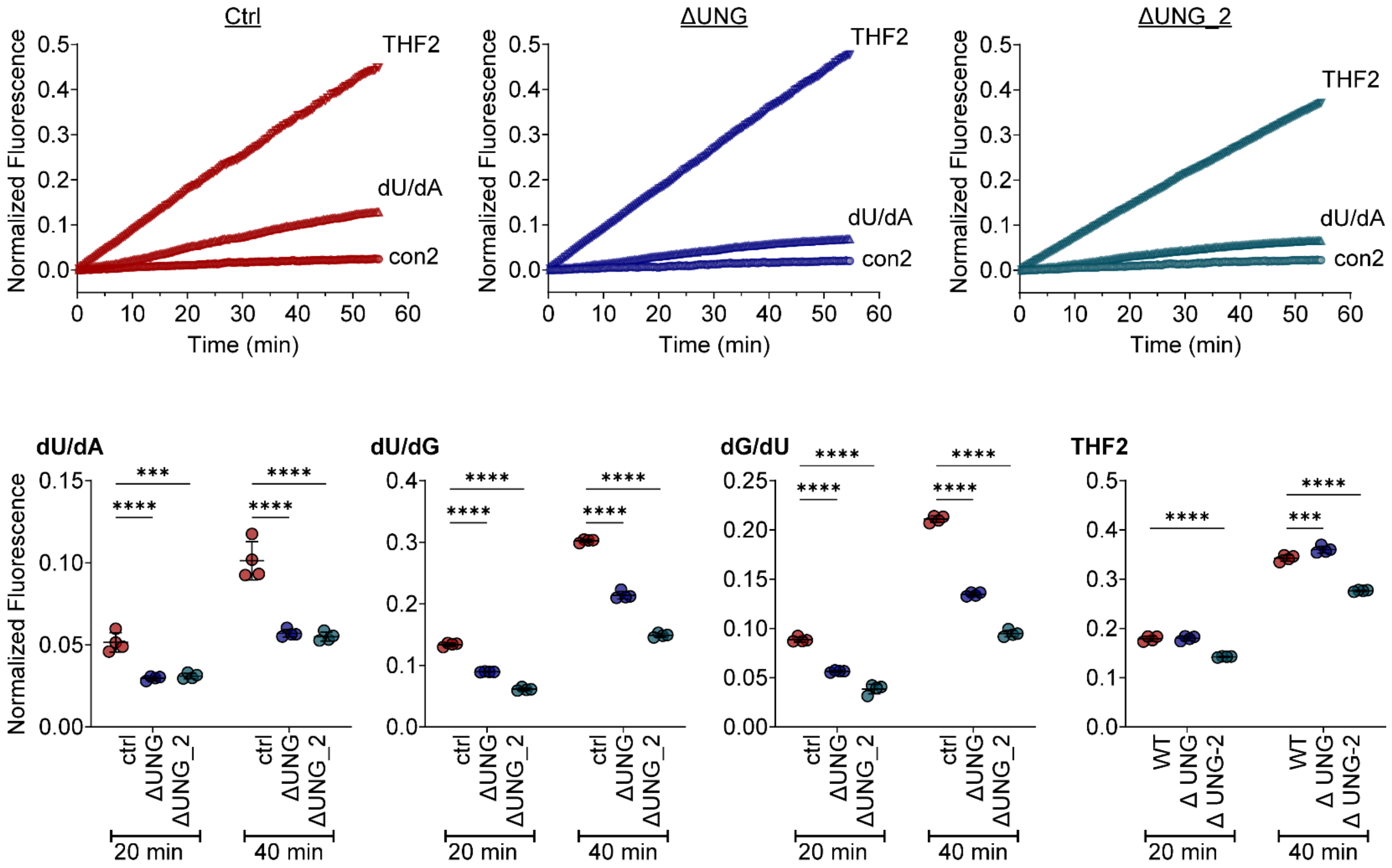


**Supplemental Figure S2: UNG deficient B16 cells have reduced glycosylase activity at dU:dA and dU:dG base pairs.**

DNA repair molecular beacons containing deoxyuridine (dU) or tetrahydrofuran (THF), which mimics an abasic sight and targets APE1, are hairpins with a 6-Fam fluorophore on the 5' end and a Dabcyl nonfluorescent quencher on the 3' end (1). The dU/dA probe contains deoxyuridine opposite adenine. The dU/dG probe contains deoxyuridine opposite guanine. The dG/dU also contains a deoxyuridine opposite guanine, but in the reverse order. UNG activity induces fluorescence since it leads to the production of an AP site, subsequently cleaved by APE1 that hydrolyzes the DNA backbone, separating the fluorophore and the quencher (2). As expected, ΔUNG and ΔUNG_2 B16 cells had significantly reduced activity against dU/dA, dU/dG, and dG/dU, compared to Cas9-expressing control cells (ctrl). As expected, ΔUNG cells did not have reduced APE1 activity, indicated by their retained reduced activity against the THF2 beacon. ΔUNG_2 cells had reduced activity against the THF2 beacon, indicating that they had reduced APE1 activity. Therefore, ΔUNG B16 cells were used throughout this study. **(Top panels)** Mean normalized fluorescence over time representing activity against the THF2, dU/dA, and negative control (con2) beacons. **(Botton panels)** Quantitation of normalized fluorescence, representing activity against the dU/dA, dU/dG, dG/dU, and THF2 beacons, at 20 and 40 minutes. n=4 replicates. Mean ± SD bars shown. ***$p<0.001$, ****$p<0.0001$ by two-way ANOVA with Sidak's multiple comparisons test.

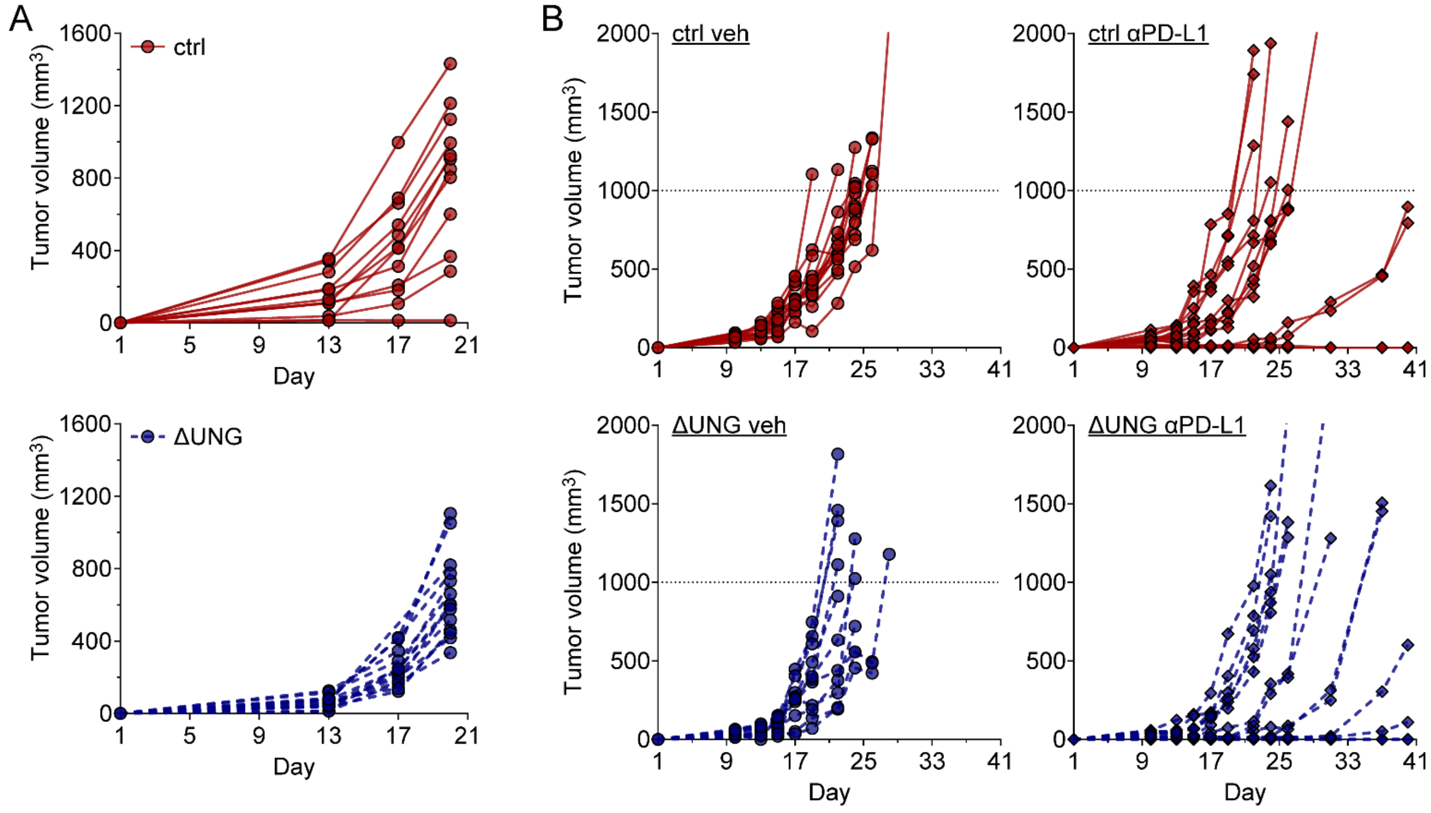


**Supplemental Figure S3. UNG deficiency alters B16 melanoma growth in immune competent mice.**

**A.** Individual tumor growth curves for Cas9-expressing control (ctrl) and UNG-deficient (ΔUNG) B16 tumors in C57BL/6 mice. Data from one experiment. n = 12 (ctrl) or 14 (ΔUNG) mice. **B.** Ctrl and ΔUNG B16 cells were injected (day 1) into C57BL/6 mice and mice were treated with 100 µg anti-PDL1 every 3 days for 6 doses, starting on day 2. Individual tumor growth curves shown until day 40, when all mice remaining mice were euthanized.

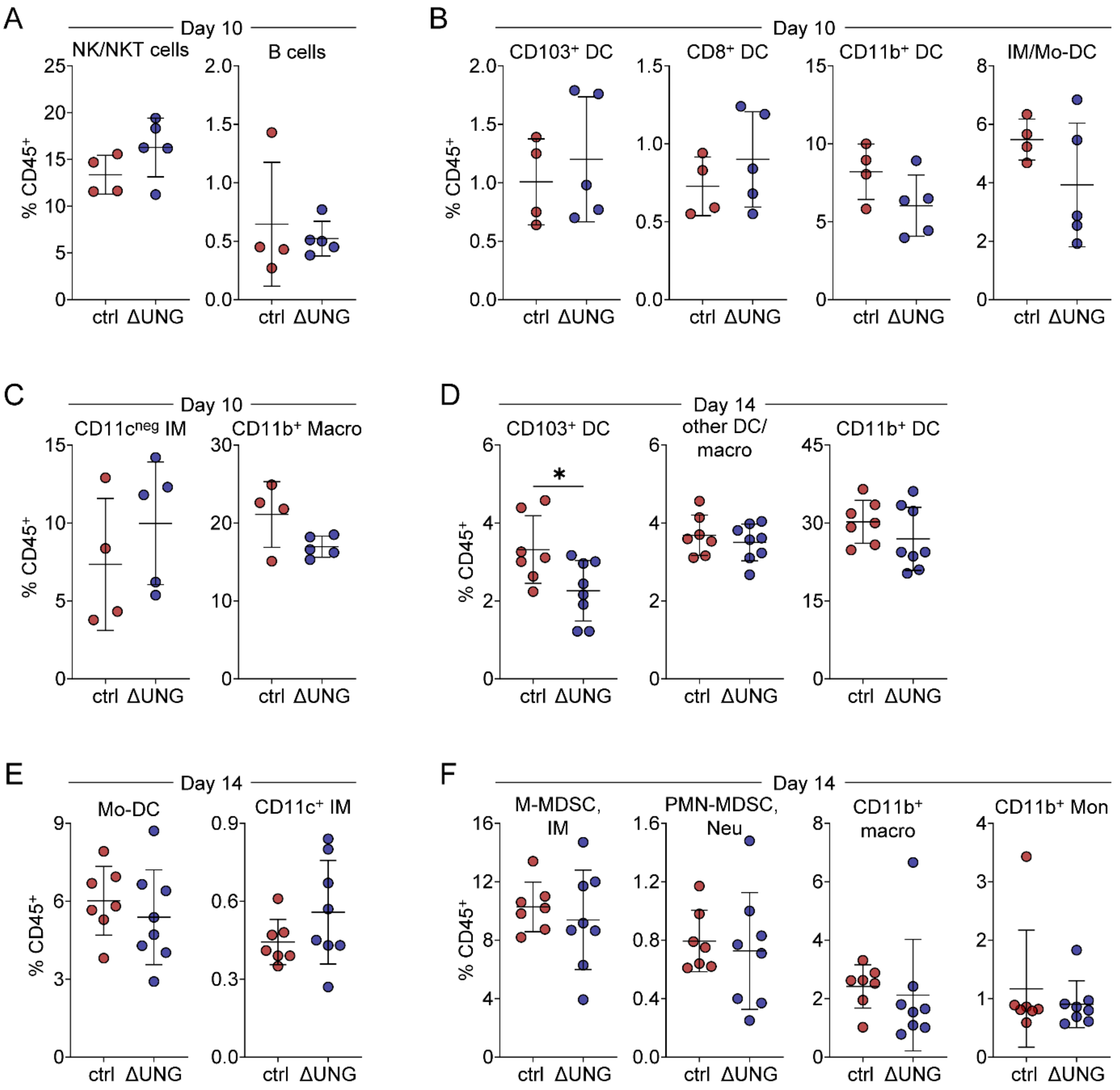


**Supplemental Figure S4. Immune profiling of control and UNG-deficient B16 tumor infiltrates**

**A-C.** Flow cytometry immunoprofiling of control (ctrl) and ΔUNG B16 tumors on days 10. Data from one experiment with n = 4 (ctrl) or 5 (ΔUNG) mice per group. **A.** Quantitation of tumor-infiltrating NK1.1$^+$ cells (NK/NKT) and B cells as percentages of the CD45$^+$ immune cell infiltrate. **B.** Quantitation of tumor-infiltrating CD11c$^+$ dendritic cell (DC) populations as percentages of the CD45$^+$ immune cell infiltrate. DC populations were subset based on expression of CD103, CD8α, CD11b, Ly-6C, and MHC-II, and include: CD103$^+$ DC, CD8$^+$ DC, CD11b$^+$ DC, and inflammatory monocyte/monocyte-derived DC (IM/Mo-DC). **C.** Quantitation of tumor-infiltrating CD11b$^+$ cell populations as percentages of the CD45$^+$ immune cell infiltrate. Subsets include C11b$^+$CD11c$^{neg}$ inflammatory monocytes (CD11c$^{neg}$ IM) and CD11b$^+$MHC-II$^+$ macrophages (CD11b$^+$ Macro). **D-F.** Flow cytometry immunoprofiling of control (ctrl) and ΔUNG B16 tumors on days 14. Data from one experiment with n = 7 (ctrl) or 8 (ΔUNG) mice per group. **D.** Quantitation of tumor-infiltrating $CD11c^{+}$ dendritic cell (DC) populations as percentages of the $CD45^{+}$ immune cell infiltrate. DC populations were first subset based on expression of CD103 and CD11b, and include: $CD103^{+}$ DC, $CD11b^{+}$ DC, and $CD11c^{+}$ cells (including $CD103^{neg}CD11b^{neg}$ DC and macrophages, other DC/macro). **E.** Quantitation of further characterized $CD11b^{+}CD11c^{+}$ dendritic cell (DC) populations as percentages of the $CD45^{+}$ immune cell infiltrate. $CD11b^{+}CD11c^{+}$ DC populations were further subdivided, based on Ly-6C and MHC-II expression, into monocyte-derived DC ($Ly\text{-}6C^{+}MHC\text{-}II^{+}$, Mo-DC) and $CD11c^{+}$ inflammatory monocytes ($Ly\text{-}6C^{+}MHC\text{-}II^{neg}$, $CD11c^{+}$ IM). **F.** Quantitation of myeloid subsets within the $CD11b^{+}CD11c^{neg}$ population as percentages of the $CD45^{+}$ immune cell infiltrate. These subsets, determined by expression of Ly-6C, Ly-6G, and MHC-II, include: $CD11b^{+}Ly\text{-}6C^{hi}Ly\text{-}6G^{neg}$ monocytic (mononuclear) myeloid-derived suppressor cells (M-MDSC) or inflammatory monocytes (IM); $CD11b^{+}Ly\text{-}6C^{int}Ly\text{-}6G^{+}$ granulocytic/polymorphonuclear myeloid-derived suppressor cells (PMN-MDSC) or neutrophils (Neu); $CD11b^{+}MHC\text{-}II^{+}$ (and Ly-6C/Ly-6G neg) macrophages ($CD11b^{+}$ macro); and $CD11b^{+}MHC\text{-}II^{neg}$ (and Ly-6C/Ly-6G neg) monocytes ($CD11b^{+}$ Mon). **A-F.** Mean ± SD bars shown. *$p<0.05$ by two-tailed, unpaired Welch's t test.

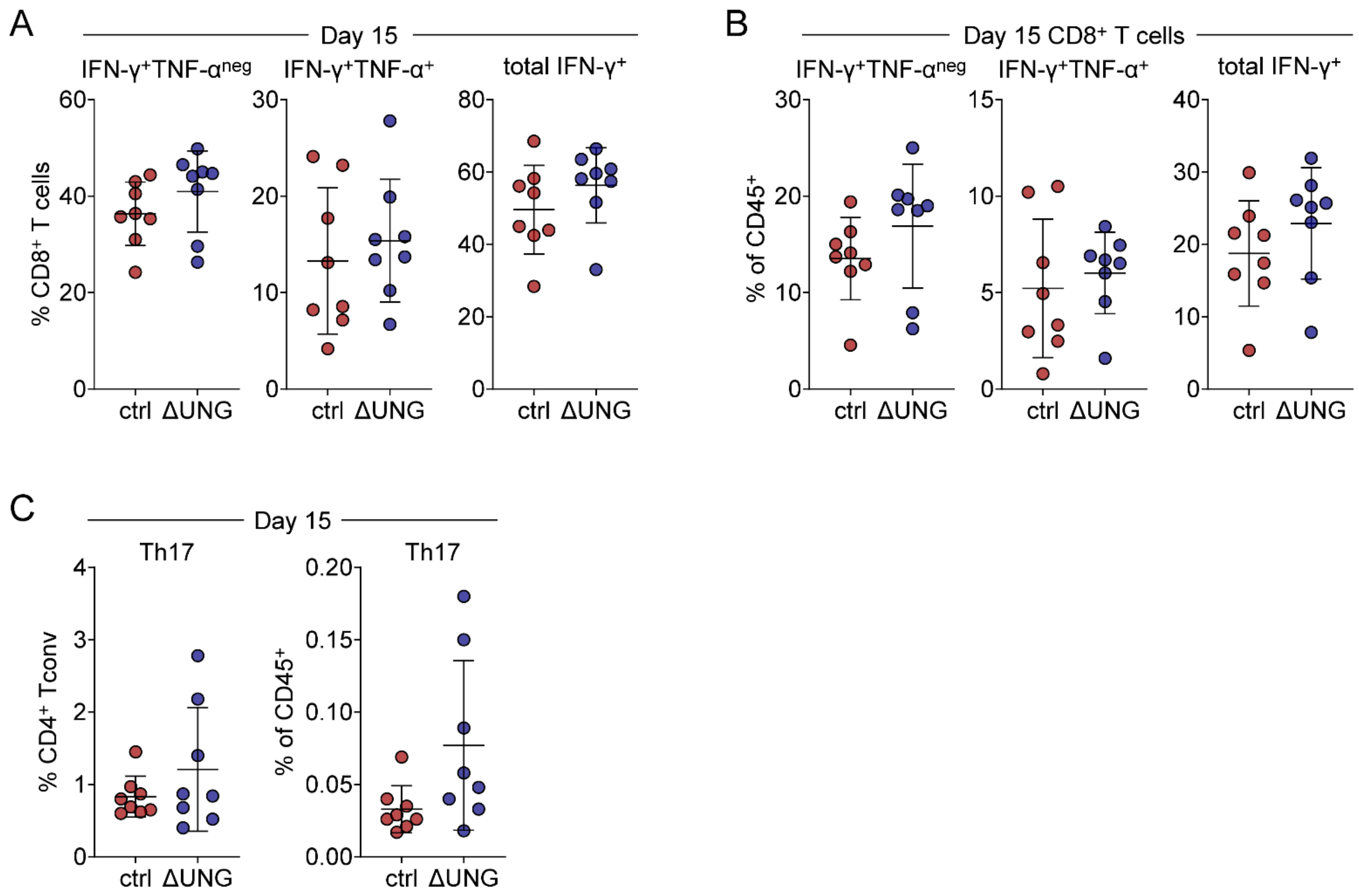


**Supplemental Figure S5. Cytokine competency of tumor-infiltrating T cells harvested from control and UNG-deficient B16 tumors on day 15.**

**A-C.** Flow cytometry analysis of cytokine-competent T cells on day 15 following *ex vivo* stimulation of control (crtl) and ΔUNG B16 tumor infiltrates with PMA/ionomycin. **A-B.** Quantitation of IFN-$\gamma^+$TNF-$\alpha^{neg}$, IFN-$\gamma^+$TNF-$\alpha^+$, and total IFN-$\gamma^+$ (TNF-$\alpha^{+/neg}$) tumor-infiltrating $CD8^+$ T cells, as percentages of the total $CD8^+$ T cell **(A)** or $CD45^+$ immune cell **(B)** infiltrate. **C.** Quantitation of IL-$17^+$ tumor-infiltrating CD4+ Tconv (Th17), as percentages of the total $CD4^+$ Tconv (left) or $CD45^+$ immune cell (right) infiltrate. **A-C.** Data from one experiment. n = 8 mice per group. Mean ± SD bars shown. No significant differences by two-tailed, unpaired Welch's t test.

A

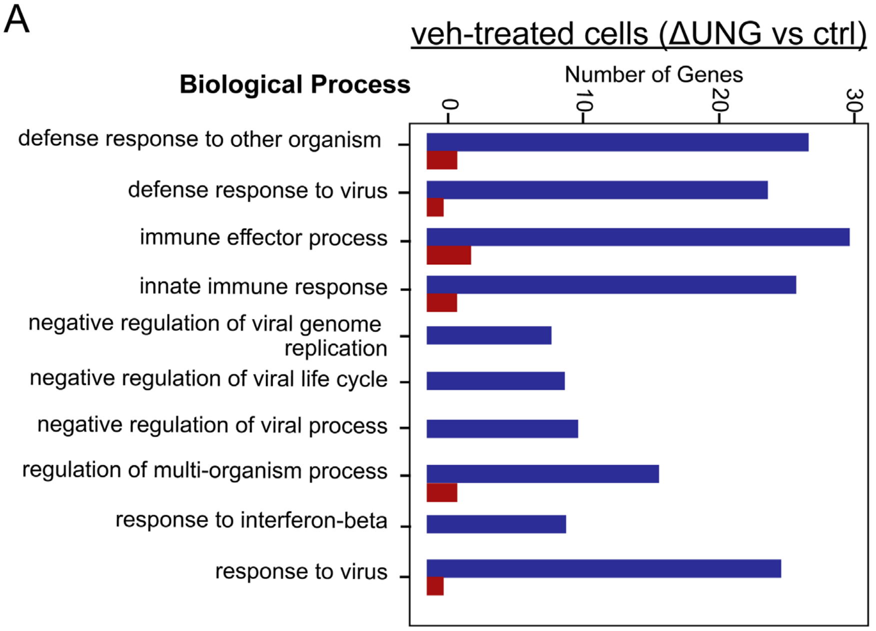


B

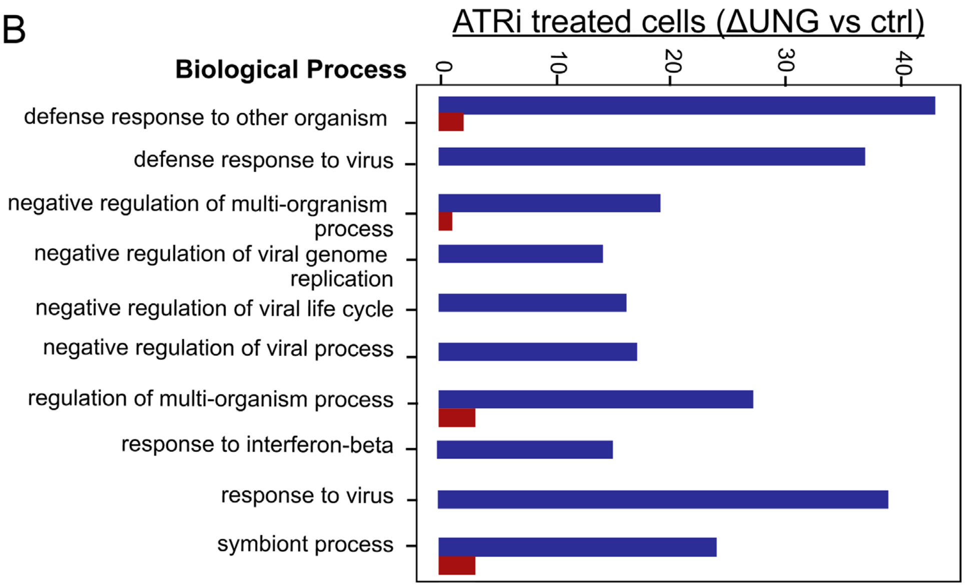


**Supplemental Figure S6. UNG deficiency represses tumor cell-intrinsic basal and ATRi-induced inflammatory signaling.**

Control (ctrl) and ΔUNG B16 cells were treated with 5 μM AZD6738 (ATRi) or vehicle (veh) for 48 h and RNA sequencing was performed with two biological replicates per condition. **A-B.** Gene Ontology analysis showing the top 10 most significantly altered biological processes in (A) veh-treated ΔUNG versus veh-treated ctrl cells, and in (B) ATRi-treated ΔUNG versus ATRi-treated ctrl cells.

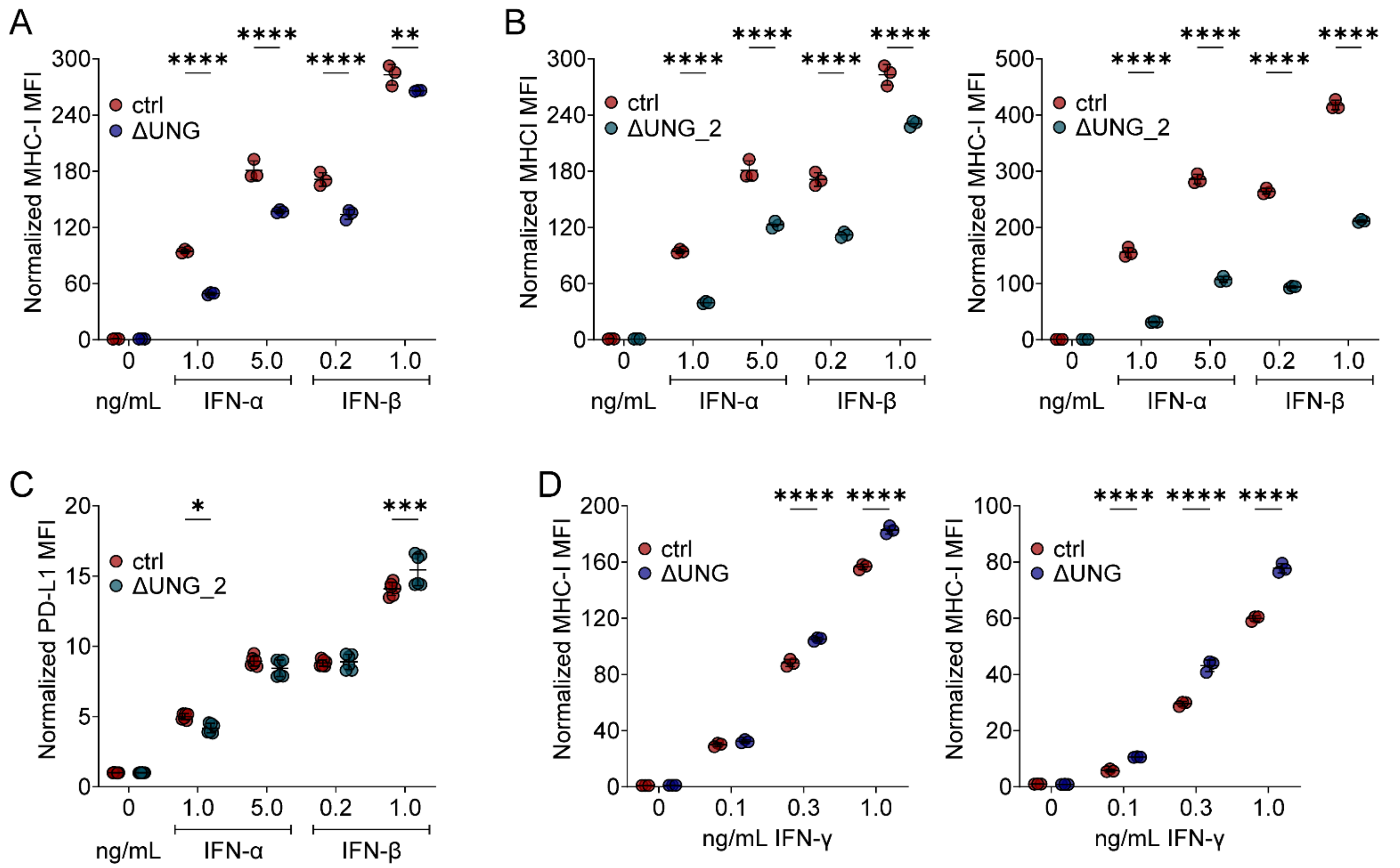


**Supplemental Figure S7. UNG deficiency alters tumor cell sensitivity to exogenous interferons *in vitro*.** **A**. MHC-I cell surface expression was analyzed on ctrl and ΔUNG B16 cells treated *in vitro* with IFN-α (1.0 or 5.0 ng/mL) or IFN-β (0.2 or 1.0 ng/mL) for 18 h. Quantitation of MHC-I MFI, normalized to the mean of untreated ctrl cells, is shown. Data from one experiment with 3 biological replicates (n=3). **B-C**. MHC-I and PD-L1 cell surface expression were analyzed on ctrl and ΔUNG clone 2 (ΔUNG _2) B16 cells treated *in vitro* with IFN-α (1.0 or 5.0 ng/mL) or IFN-β (0.2 or 1.0 ng/mL) for 18 h. **B.** Quantitation of MHC-I MFI, normalized to the mean of untreated ctrl cells, is shown for two independent experiments, each with 3 biological replicates (n=3 per experiment). **C.** Quantitation of PD-L1 MFI, normalized to the mean of untreated ctrl cells, combined from two independent experiments, each with 3 biological replicates (n=9 in total). **D**. MHC-I cell surface expression was analyzed on ctrl and ΔUNG B16 cells treated *in vitro* with IFN-γ (0.1, 0.3, or 1.0 ng/mL) for 18 h. Quantitation of MHC-I MFI, normalized to the mean of untreated ctrl cells, is shown for two independent experiments, each with 3 biological replicates (n=3 per experiment). **A-D.** Mean ± SD bars shown. *p<0.05, **p<0.01, ***p<0.001, ****p<0.0001 by two-way ANOVA with Sidak's **(A-C)** or Tukey's **(D)** multiple comparisons test.

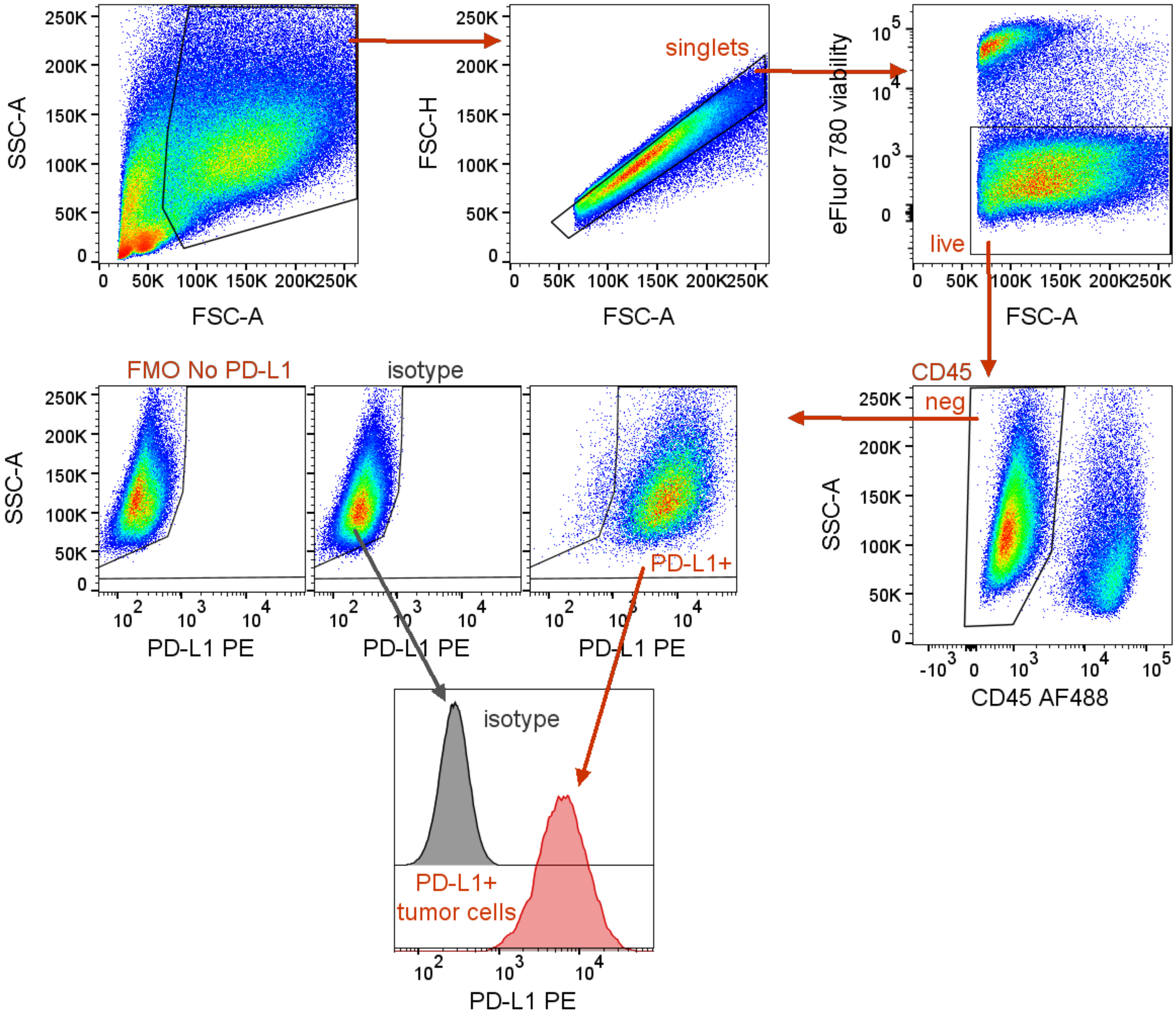


**Supplemental Figure S8. Gating strategy for day 15 profiling of tumor PD-L1 expression *in vivo*.**

Any unstable portions of the run were excluded, and cells were gated based on FSC-A vs. SSC-A, followed by gating on FSC-H vs. FSC-A (doublet exclusion). Live (viability dye negative), $CD45^{neg}$ tumor cells were gated based on PD-L1 expression (gate determined by fluorescence minus one (FMO) control without PD-L1 PE) and median fluorescence intensities (MFI) were determined. The histogram shows PD-L1 staining intensity for PD-$L1^{+}$ cells in a representative tumor versus the background signal for isotype control staining on total $CD45^{neg}$ tumor population.

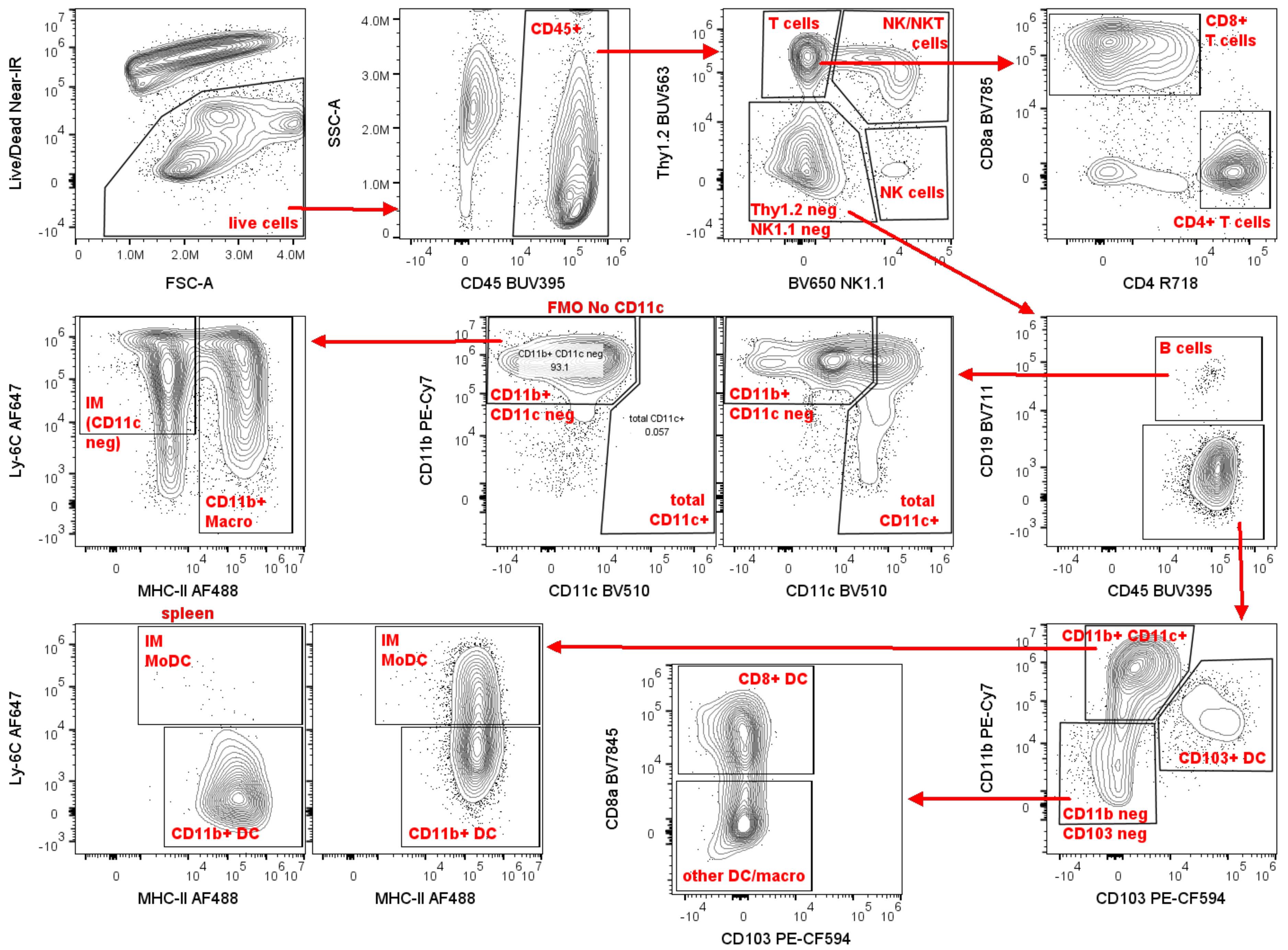
Live/Dead Near-IR
FSC-A
live cells
SSC-A
CD45 BUV395
CD45+
Thy1.2 BUV563
BV650 NK1.1
T cells
NK/NKT cells
NK cells
Thy1.2 neg NK1.1 neg
CD8a BV785
CD4 R718
CD8+ T cells
CD4+ T cells
FMO No CD11c
CD11b PE-Cy7
CD11c BV510
CD11b+ CD11c neg
93.1
total CD11c+
0.057
CD11b+ CD11c neg
total CD11c+
CD19 BV711
CD45 BUV395
B cells
Ly-6C AF647
MHC-II AF488
IM (CD11c neg)
CD11b+ Macro
spleen
IM MoDC
CD11b+ DC
CD8a BV7845
CD103 PE-CF594
CD8+ DC
other DC/macro
CD11b+ CD11c+
CD103+ DC
CD11b neg CD103 neg

**Supplemental Figure S9. Gating strategy for day 10 profiling of tumor-infiltrating immune cells using spectral flow cytometry.**

Any unstable portions of the run were excluded, and cells were gated based on FSC-A vs. SSC-A, followed by gating on FSC-H vs. FSC-A (doublet exclusion). Live cells were identified as Live/Dead Near-IR viability dye negative. $CD45^{+}$ immune cells were gated. $CD45^{+}$ cells negative for the lineage markers Thy1.2 and NK1.1 were gated to subset T cells, $Thy1.2^{+}$ NK/NKT cells, and $Thy1.2^{neg}$ NK cells. $Thy1.2^{neg}NK1.1^{neg}$ cells were subset into B cells ($CD19^{+}$) and non-lymphocytes (Thy1.2/NK1.1/CD19 negative).

Non-lymphocytes were subdivided based on expression of CD11b and CD11c into $CD11b^{+}CD11c^{neg}$ cells and total $CD11c^{+}$ cells. $CD11b^{+}CD11c^{neg}$ cells were subdivided based expression of Ly-6C and MHC-II into inflammatory monocytes (IM, $CD11b^{+}Ly\text{-}6C^{+}CD11c^{neg}MHC\text{-}II^{neg}$) or $CD11b^{+}MHC\text{-}II^{+}$ macrophages ($CD11b^{+}$ Macro).

Total $CD11c^{+}$ cells were subdivided based on expression of CD103 and CD11b. $CD11c^{+}CD103^{+}CD11b^{neg}$ cells were defined as $CD103^{+}$ dendritic cells ($CD103^{+}$ DC), while $CD11c^{+}CD103^{neg}CD11b^{neg}$ were subset based on CD8α expression into $CD8^{+}$ DC ($CD11c^{+}CD8\alpha^{+}$ and CD103/CD11b negative) and other DC/macro ($CD11c^{+}$ and CD8α/CD103/CD11b negative).

$CD11c^{+}CD11b^{+}CD103^{neg}$ cells were subdivided based on Ly-6C and MHC-II expression into inflammatory monocytes/monocyte-derived dendritic cells (IM/Mo-DC, $CD11c^{+}CD11b^{+}Ly\text{-}6C^{+}MHC\text{-}II^{+}$) and $CD11b^{+}$ dendritic cells ($CD11b^{+}$ DC, $CD11c^{+}CD11b^{+}Ly\text{-}6C^{neg}MHC\text{-}II^{+}$).

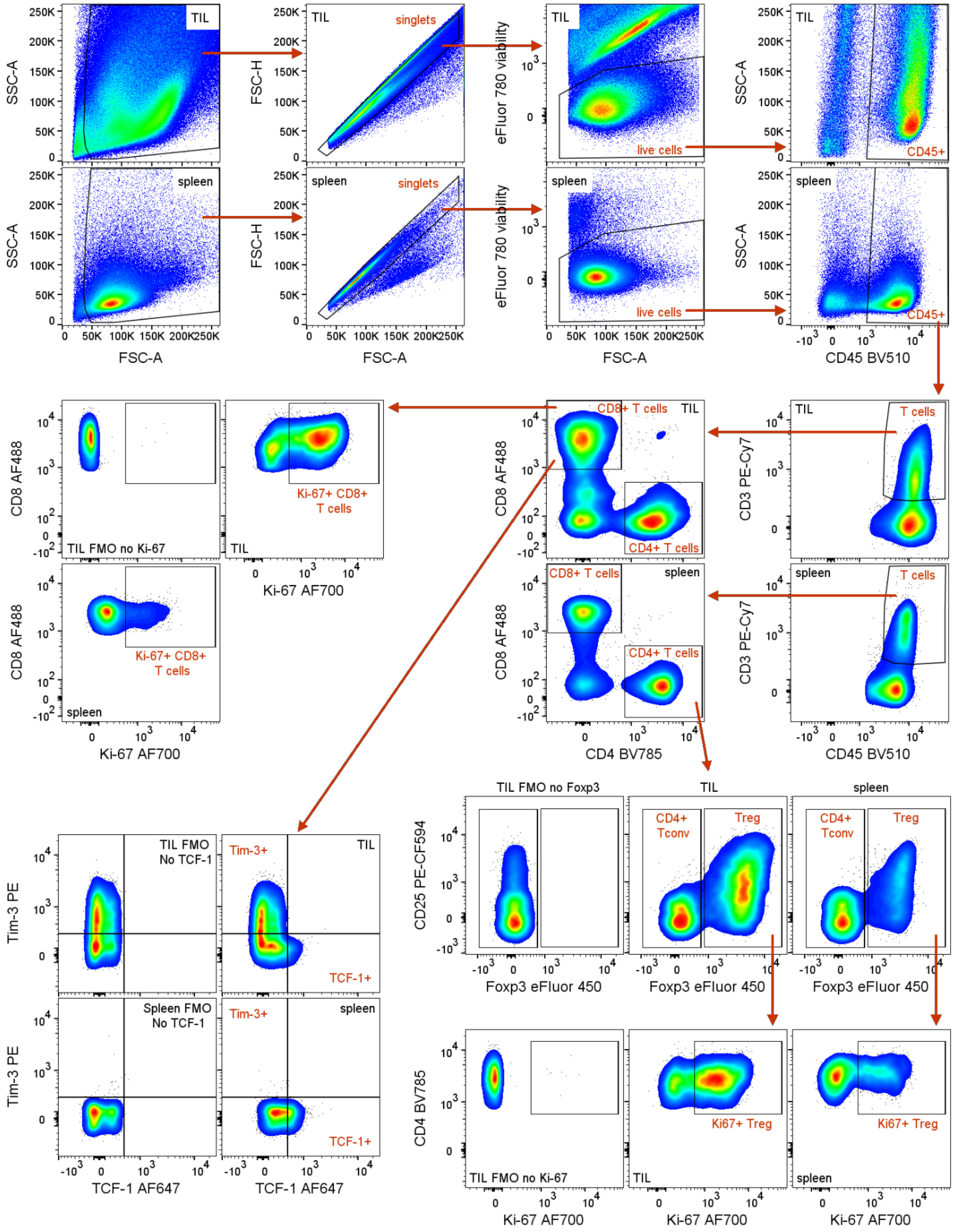


(same gating strategy for Ki-67 in CD4+ Tconv)

**Supplemental Figure S10. Gating strategy for day 14 profiling of tumor-infiltrating T cells.**

Any unstable portions of the run were excluded, and cells were gated based on FSC-A vs. SSC-A, followed by gating on FSC-H vs. FSC-A (doublet exclusion). Live (viability dye negative) cells were gated on CD45 expression, and $CD45^+$ immune cells were gated based expression CD3 to identify $CD3^+$ T cells, which were further subset into $CD8^+$ and $CD4^+$ T cells. $CD4^+$ T cells were further divided, based on expression of Foxp3 and CD25, into $Foxp3^{neg}$ conventional $CD4^+$ T cells ($CD4^+$ Tconv) and $Foxp3^+$ regulatory T cells (Treg). The marker CD25 was included to aid in resolution of the $CD4^+$ Tconv and Treg populations. TIL and spleen fluorescence minus one (FMO) controls (without Foxp3 eFluor 450) were included to determine gating for Foxp3. The $CD8^+$ T cell, $CD4^+$ Tconv, and Treg populations were all analyzed for expression of Ki-67. Gating for Ki-67 was determined using the spleen gating control sample in addition to FMO controls (without Ki-67 AF700). $CD8^+$ T cells were also examined for expression of Tim-3 and TCF-1, with gating determined using TIL and spleen FMO controls (without TCF-1 AF647) for the TCF-1 gating and the spleen gating control for the Tim-3 gate. Since no differences in Tim-3 expression across groups were observed, quantitation was not included in the manuscript.

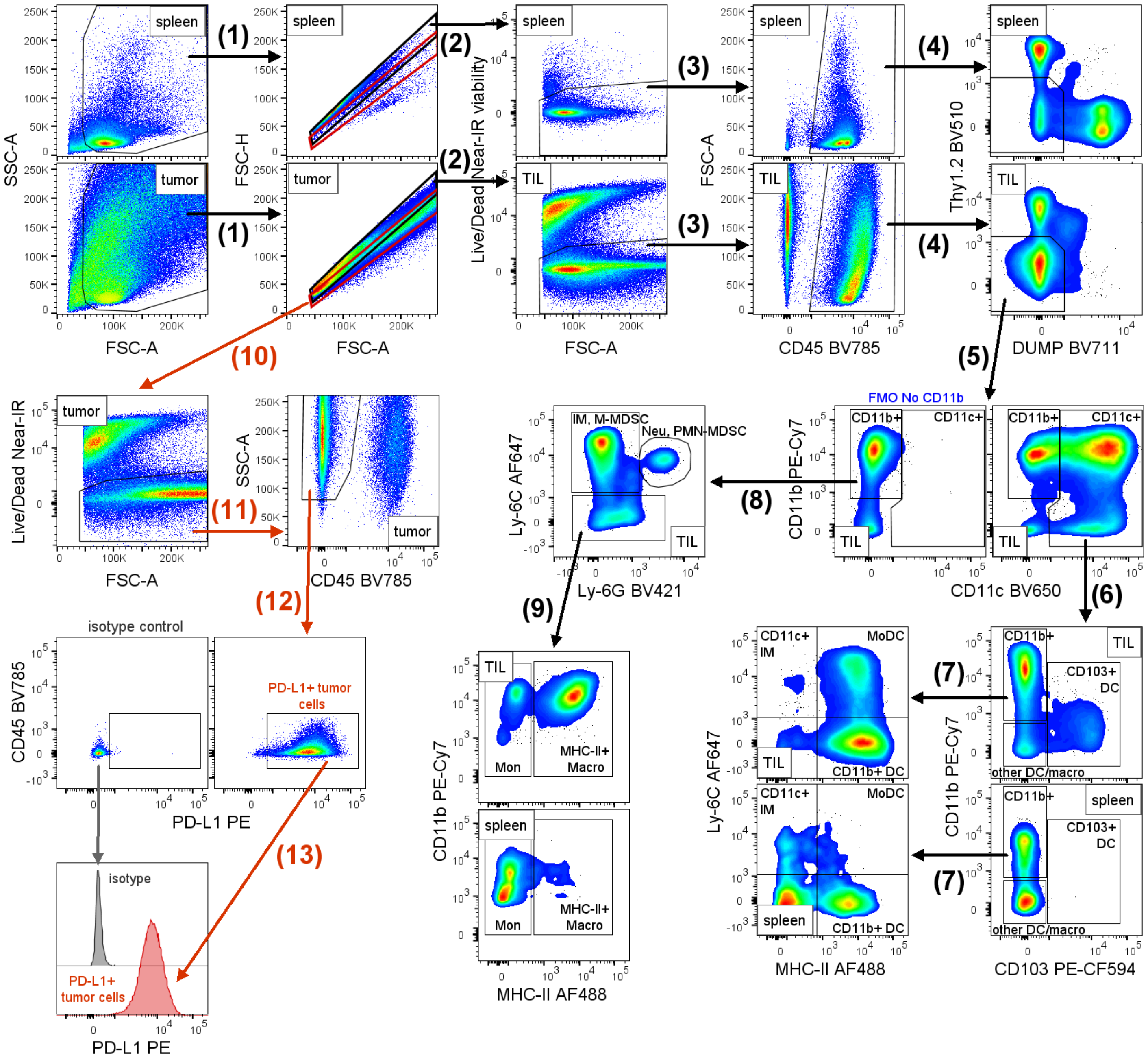


**Supplemental Figure S11. Gating strategy for day 14 profiling of tumor-infiltrating myeloid cells and tumor PDL1 expression.**

Any unstable portions of the run were excluded, and cells were gated based on FSC-A vs. SSC-A, followed by **(1)** gating on FSC-H vs. FSC-A (doublet exclusion). Since immune cells and tumor cells have different FSC-H vs. FSC-A characteristics, two single cell gates were created. The **black** gate was drawn based on the spleen sample to identify the single cell immune population for myeloid profiling. The **red** gate was drawn to identify the single cell tumor cell population for analysis of tumor PD-L1 expression.

Immune cells within the **black single cell gate** were subset into myeloid populations. **(2)** Live cells were identified as Live/Dead Near-IR viability dye negative. CD45⁺ immune cells were gated. **(4)** CD45⁺ cells negative for the lineage markers Thy1.2, CD19, and NK1.1 were gated to remove T cells, B cells and NK cells, respectively

(CD19 and NK1.1 antibodies were combined in a BV711 DUMP channel). **(5)** Lineage negative cells were subset into total CD11c$^+$ (CD11b$^+$ or CD11b$^{neg}$) cells and CD11b$^+$CD11c$^{neg}$ cells. **(6)** Total CD11c$^+$ cells were subdivided based on expression of CD103 and CD11b. CD11c$^+$CD103$^+$ cells were defined as CD103$^+$ dendritic cells (CD103$^+$ DC), while CD11c$^+$CD103$^{neg}$CD11b$^{neg}$ were defined as other dendritic cells or macrophages (other DC/macro) without further characterization. **(7)** CD11c$^+$CD11b$^+$CD103$^{neg}$ cells were subdivided based on Ly-6C and MHC-II expression as follows: CD11c$^+$CD11b$^+$Ly-6C$^{neg}$MHC-II$^+$ cells were defined as CD11b$^+$ dendritic cells (CD11b$^+$ DC), CD11c$^+$CD11b$^+$Ly-6C$^+$MHC-II$^{neg}$ cells were defined as CD11c$^+$ inflammatory monocytes (CD11c$^+$ IM), and CD11c$^+$CD11b$^+$Ly-6C$^+$MHC-II$^+$ cells were defined as monocyte-derived dendritic cells (Mo-DC). **(8)** Lineage negative CD11b$^+$CD11c$^{neg}$ cells were subdivided based expression of Ly-6C and Ly-6C. CD11b$^+$CD11c$^{neg}$Ly-6C$^{hi}$Ly-6G$^{neg}$ were defined as inflammatory monocytes (IM) or monocytic (mononuclear) myeloid-derived suppressor cells (M-MDSC). CD11b$^+$CD11c$^{neg}$Ly-6C$^{int}$Ly-6G$^+$ were defined as neutrophils (Neu) or granulocytic (polymorphonuclear) myeloid-derived suppressor cells (PMN-MDSC). **(9)** Cells negative for both Ly-6C and Ly-6G were subset based on MHC-II expression. CD11b$^+$MHC-II$^{neg}$ (and CD11c/Ly-6C/Ly-6G negative) cells were defined as monocytes (CD11b$^+$MHC-II$^{neg}$ Mon), while CD11b$^+$MHC-II$^+$ (and CD11c/Ly-6C/Ly-6G negative) were defined as MHC-II$^+$ macrophages (CD11b$^+$MHC-II$^+$ Macro).

Within the **red single cell gate**, **(10)** live cells were identified as Live/Dead Near-IR viability dye negative. **(11)** CD45$^{neg}$ tumor cells were gated. **(12)** CD45$^{neg}$ tumor cells were then gated based expression of PD-L1 (determined empirically using a fluorescence minus one (FMO) control without PD-L1 PE (not shown), and median fluorescence intensities (MFI) were determined. An isotype control tumor sample was included to verify minimal background signal. **(13)** The histogram depicts PD-L1 staining intensity for the PD-L1+ population in a representative tumor versus background signal for isotype control staining on total CD45$^{neg}$ tumor population.

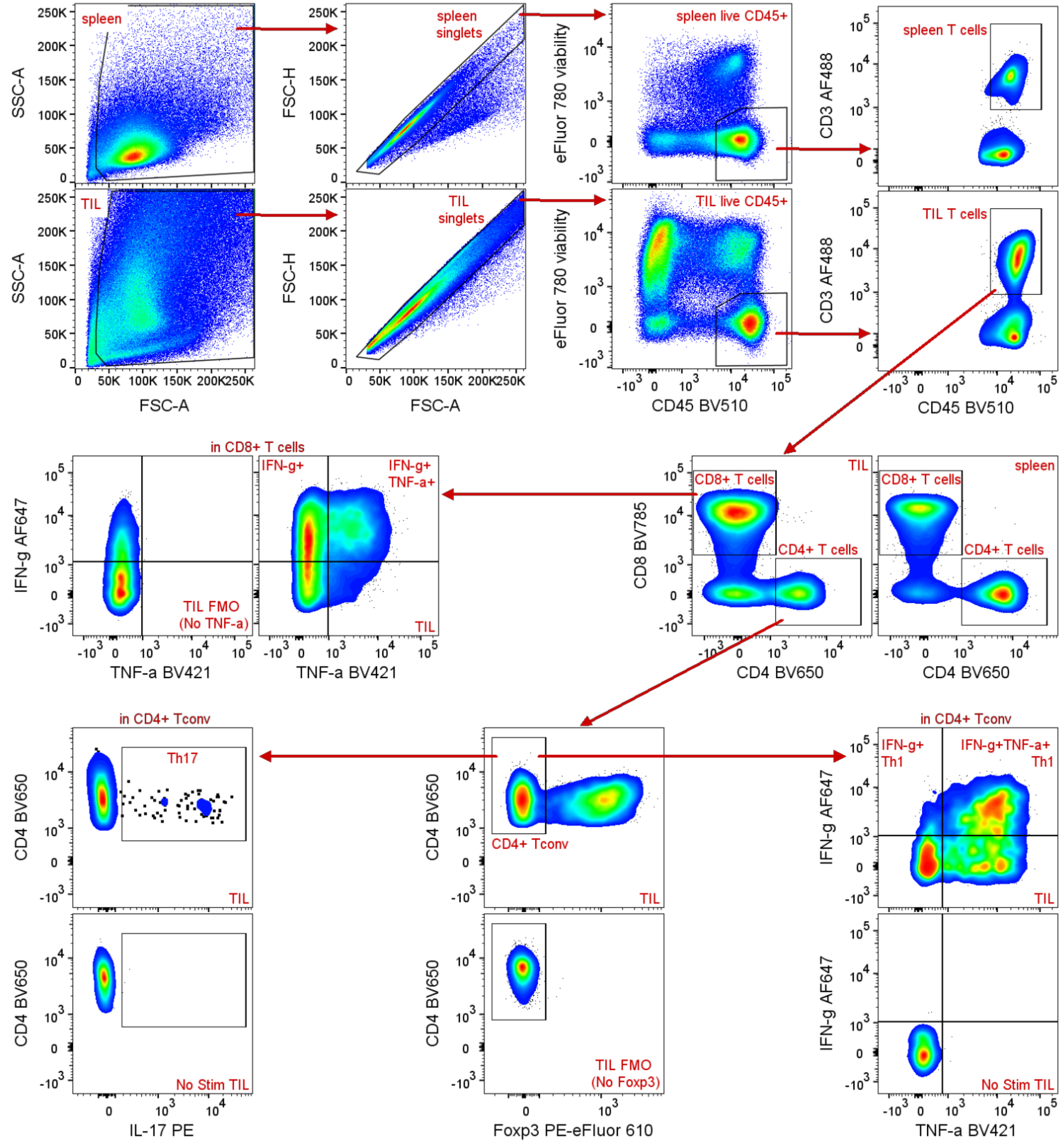


**Supplemental Figure S12. Gating strategy for day 15 profiling of tumor-infiltrating, cytokine-competent T cells.**

Any unstable portions of the run were excluded, and cells were gated based on FSC-A vs. SSC-A, followed by gating on FSC-H vs. FSC-A (doublet exclusion). Live (viability dye negative), $CD45^{+}$ immune cells were gated, and total T cells ($CD3^{+}$) were divided into $CD8^{+}$ and $CD4^{+}$ T cell populations. $CD8^{+}$ T cells were examined for production of IFN-γ and TNF-α. $CD4^{+}$ T cells were examined for Foxp3 expression to exclude regulatory T cells from the conventional CD4⁺ T cells (CD4⁺ Tconv), which were examined for production of IFN-γ and TNF-α (to identify the Th1 subset) or IL-17 (to identify the Th17 subset). Unstimulated TIL and unstimulated spleen (No Stim) controls and No TNF-α fluorescence minus one (FMO) controls were used to determine gating for IFN-γ and TNF-α in both CD8⁺ T cells and CD4+ Tconv. No Foxp3 FMO controls were used to determine gating for CD4⁺ Tconv (ie. Foxp3$^{neg}$). IL-17 gating was determined using the No Stim controls.

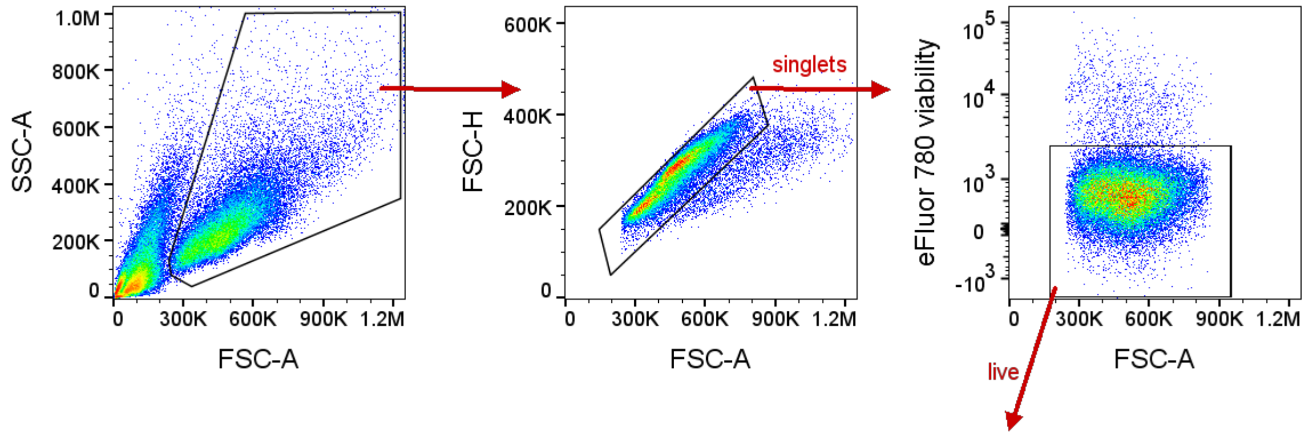


**Supplemental Figure S13. Gating strategy for MHC-I and PD-L1 cell surface expression *in vitro.***
Cells were gated based on FSC-A vs. SSC-A, followed by gating on FSC-H vs. FSC-A (doublet exclusion). Live cells (eFluor 780 viability dye negative) were analyzed for the relative surface expression of MHC-I and PD-L1, as measured by their median fluorescence intensity (MFI).

**Resources and Reagents**

| Plasmids for lentivirus production and transduction | Source | Identifier |
|---|---|---|
| pMDLg/pRRE | Addgene | 12251 |
| pRSV-Rev | Addgene | 12253 |
| pMD2.G | Addgene | 12259 |
| pLENTI-CRISPR-V2-control gRNA_1 | This study | Sobol Lab 2014 |
| pLENTI-CRISPR-V2-control gRNA_2 | This study | Sobol Lab 2015 |
| pLV-hCas9-T2A-Puro-U6-mUng-gRNA9 | This study | Sobol Lab 2416 |

| *ung* Primers used for Sanger Sequencing | Source |
|---|---|
| Pair 1 Forward: GTTAAATCAGGTTGGCGGGCTG | IDT |
| Pair 1 Reverse: GGCCATCCCTTCTAGTCTGTGC | IDT |
| Pair 2 Forward: GAAGTCAGGAAAGACGATCGAGGC | IDT |
| Pair 2 Reverse: CGTTGCCTACACTCAATAGACTGAG | IDT |

| Oligonucleotides/gRNA for Molecular Beacon Assay | Source | Lesion |
|---|---|---|
| 6-Fam-CCA CTA TTG AAT TGA CAC GCC ATG TCG ATC AAT TCA ATA GTG G-Dabcyl | IDT | FD-Con2[7] |
| 6-Fam-CCA CTX TTG AAT TGA CAC GCC ATG TCG ATC AAT TCA ATA GTG G-Dabcyl | IDT | FD-THF2, where X indicates the location of lesion[7] |
| 6-Fam-CCA CTX TTG AAT TGA CAC GCC ATG TCG ATC AAT TCA AAA GTG G-Dabcyl | IDT | FD-dU/A, where X indicates the location of the lesion[7] |
| 6-Fam-CCA CTX GTG AAT TGA CAG CCC ATG TGC ATC AAT TCA CGA GTG G-Dabcyl | IDT | FDB-dU/G, where X indicates the location of the lesion[7] |
| 6-Fam-CCA CCG TTG AAT TGA CAG CCC ATG TGC ATC AAT TCA AXG GTG G-Dabcyl | IDT | FDB-G/dU, where X indicates the location of the lesion[7] |

| Flow Cytometry | Source | Identifier |
|---|---|---|
| AF647 anti-mouse TCF-7/TCF-1, clone S33-966 (1:200 Dilution) | BD Biosciences | 566693 |
| BUV395 anti-mouse CD45, clone 30-F11 (1:250 Dilution) | BD Biosciences | 564279 |
| BUV563 anti-mouse CD90.2 (Thy1.2), clone 53-2.1 (1:400 Dilution) | BD Biosciences | 741213 |
| BV421 anti-mouse CD274 (PD-L1), clone 10F.9G2 (1:100 Dilution) | BD Biosciences | 568309 |
| BV421 anti-mouse H-2Ld/H-2Db (MHC-I), clone 28-14-8 (1:100 Dilution) | BD Biosciences | 742468 |
| BV421 rat IgG2b, κ isotype control, clone R35-38 (1:100 Dilution) | BD Biosciences | 562603 |
| BV650 anti-mouse CD4, clone GK1.5 (1:500 Dilution) | BD Biosciences | 563747 |
| PE-CF594 anti-mouse CD25, clone PC61 (1:250 Dilution) | BD Biosciences | 562695 |
| PE-CF594 anti-mouse CD103, clone M290 (1:250 Dilution) | BD Biosciences | 565849 |
| R718 anti-mouse CD4, clone RM4-5 (1:600 Dilution) | BD Biosciences | 566939 |
| AF488 anti-mouse CD3, clone 17A2 (1:250 Dilution) | BioLegend | 100212 |
| AF488 anti-mouse CD8α, clone 53-6.7 (1:250 Dilution) | BioLegend | 100723 |
| AF488 anti-mouse CD45, clone 30-F11 (1:250 Dilution) | BioLegend | 103122 |

| | | |
|---|---|---|
| AF488 anti-mouse I-A/I-E (MHC-II), clone M5/114.15.2 (1:500 Dilution) | BioLegend | 107615 |
| AF647 anti-mouse IFN-γ (clone XMG1.2) (1:500 Dilution) | BioLegend | 505816 |
| AF647 anti-mouse Ly-6C, clone HK1.4 (1:500 Dilution) | BioLegend | 128010 |
| AF700 anti-mouse Ki67, clone 16A8 (1:200 Dilution) | BioLegend | 652420 |
| BV421 anti-mouse Ly-6G, clone 1A8 (1:250 Dilution) | BioLegend | 127628 |
| BV421 anti-mouse TNF-α, clone MP6-XT22 (1:250 Dilution) | BioLegend | 506328 |
| BV510 anti-mouse CD11c, clone N418 (1:100 Dilution) | BioLegend | 117353 |
| BV510 anti-mouse CD45, clone 30-F11 (1:250 Dilution) | BioLegend | 103138 |
| BV510 anti-mouse Thy1.2 (CD90.2), clone 30-H12 (1:250 Dilution) | BioLegend | 105335 |
| BV650 anti-mouse CD11c, clone N418 (1:100 Dilution) | BioLegend | 117339 |
| BV650 anti-mouse NK-1.1, clone PK136 (1:250 Dilution) | BioLegend | 108736 |
| BV711 anti-mouse CD19, clone 6D5 (1:250 Dilution) | BioLegend | 115555 |
| BV711 anti-mouse NK-1.1, clone PK136 (1:250 Dilution) | BioLegend | 108745 |
| BV785 anti-mouse CD4, clone GK1.5 (1:500 Dilution) | BioLegend | 100453 |
| BV785 anti-mouse CD8α, clone 53-6.7 (1:250 Dilution) | BioLegend | 100750 |
| BV785 anti-mouse CD45, clone 30-F11 (1:250 Dilution) | BioLegend | 103149 |
| PE anti-mouse IL-17A, clone TC11-18H10.1 (1:200 Dilution) | BioLegend | 506903 |
| PE-Cy7 anti-mouse/human CD11b, clone M1/70 (1:250 Dilution) | BioLegend | 101215 |
| PE anti-mouse CD274 (PD-L1), clone 10F.9G2 (1:100 Dilution *in vitro*, 1:250 Dilution *in vivo*) | BioLegend | 124308 |
| PE anti-mouse CD366 (Tim-3), clone RMT3-23 (1:250 Dilution) | BioLegend | 119703 |
| PE anti-mouse H-2Ld/H-2Db (MHC-I), clone 28-14-8 (1:100 Dilution) | BioLegend | 114507 |
| PE mouse IgG2a, κ isotype control, clone MOPC-173 (1:100 Dilution) | BioLegend | 400211 |
| PE rat IgG2b, κ isotype control, RTK4530 (1:100 Dilution *in vitro*, 1:250 Dilution *in vivo*) | BioLegend | 400607 |
| PE-Cy7 anti-mouse CD3, clone 17A2 (1:250 Dilution) | BioLegend | 100219 |
| Per-CP anti-mouse Ly-6G, clone 1A8 (1:250 Dilution) | BioLegend | 127653 |
| eFluor 450 anti-mouse/rat Foxp3, clone FJK-16s (1:200 Dilution) | Invitrogen | 48-5773-82 |

| | | |
|---|---|---|
| PE-eFluor 610 anti-mouse/rat Foxp3, clone FJK-16s (1:200 Dilution) | Invitrogen | 61-5773-80 |
| BD Horizon™ Brilliant Stain Buffer Plus (1:5 Dilution) | BD Biosciences | 566385 |
| TruStain FcX™ PLUS (anti-mouse CD16/32) Antibody (1:100 Dilution) | BioLegend | 156604 |
| True-Stain Monocyte Blocker™ (1:20 Dilution) | BioLegend | 426103 |
| FluoroFix buffer | BioLegend | 422101 |
| eBioscience Protein Transport Inhibitor Cocktail (500X) | Invitrogen | 00-4980-93 |
| eBioscience Cell Stimulation Cocktail (plus protein transport inhibitors) (500X) | Invitrogen | 00-4975-03 |
| eBioscience fixable viability dye eFluor 780 (1:2000-1:3000 Dilution) | Invitrogen | 65-0865-18 |
| LIVE/DEAD™ Fixable Near-IR dead cell stain (1:1000 Dilution) | Invitrogen | L10119 |
| eBioscience flow cytometry staining (FCS) buffer | Invitrogen | 00-4222-26 |
| eBioscience Permeabilization Buffer (10X) | Invitrogen | 00-8333-56 |
| eBioscience Fixation/Permeabilization Concentrate | Invitrogen | 00-5123-43 |
| eBioscience Fixation/Permeabilization Diluent | Invitrogen | 00-5223-56 |
| eBioscience normal mouse serum (used at 5% in 1x PBS) | Invitrogen | 24-5544-94 |
| OneComp eBeads compensation beads | Invitrogen | 01-1111-42 |
| UltraComp eBeads Plus compensation beads | Invitrogen | 01-3333-42 |
| Collagenase Type IV, Filtered | Worthington | LS004209 |
| Deoxyribonuclease I | Worthington | LS002139 |
| Trypsin Inhibitor, Soybean (Animal-Free) | Worthington | LS003587 |
| **Chemicals/Inhibitors/Reagents** | **Source** | **Identifier** |
| AZD6738 (ATRi) | AstraZeneca | |
| Recombinant mouse IFN-α1 | BioLegend | 751802 |
| Recombinant mouse IFN-β1 | BioLegend | 581302 |
| Recombinant Murine IFN-γ (Animal-Free) | Peprotech | AF-315-05 |
| inVivoPlus anti-mouse PD-L1 (B7-H1) | BioXCell | BP0101 |
| inVivoPure pH 6.5 Dilution Buffer | BioXCell | IP0065 |
| Puromycin | InvivoGen | ant-pr-1 |
| DMEM | Lonza | 12-604F |
| Fetal Bovine Serum (FBS) | GEMINI Bioproducts | 900-108 |
| Pen/Strep | Gibco | 15140-122 |
| 0.05% Trypsin-EDTA (1x) | Gibco | 25300-054 |
| StemPro™ Accutase™ Cell Dissociation Reagent | Gibco | A1110501 |
| Phosphate buffered saline (PBS) | Fisher | BP2944-100 |
| Trypan Blue Solution (0.4%) | Millipore Sigma | T8154 |
| Dimethyl sulfoxide (DMSO) | Millipore Sigma | D2438-5X10ML |
| Propylene Glycol | Millipore Sigma | P4347 |
| Trizol | Ambion | 15596026 |
| Ethanol | Decon Labs | 2716 |
| ProcartaPlex Cell Lysis Buffer | Invitrogen | EPX-99999-000 |
| PMSF (phenylmethanesulfonyl fluoride) | ThermoFisher Scientific | 36978 |
| Tween-20 | Fisher | BP337-500 |

| | | |
|---|---|---|
| EDTA (0.5 M, pH 8.0) (for erythrocyte lysis buffer) | Fisher | BP2482-500 |
| Ammonium Chloride (for erythrocyte lysis buffer) | Fisher | A661-500 |
| Sodium Bicarbonate (for erythrocyte lysis buffer) | Fisher | BP328-500 |
| Glycerol | EMD | GX0185-5 |
| EDTA (0.5, pH 8.0) | Boston Bio Products | BM-150 |
| **Commercial Kits** | **Source** | **Identifier** |
| TransIT-X2 Dynamic Delivery System | Mirus Bio | MIR 6003 |
| Lenti-X Concentrator | Takara Bio | 631231 |
| PureLink Genomic DNA Mini Kit | Invitrogen | K182002 |
| GeneJET PCR Purification Kit | ThermoFisher Scientific | K0702 |
| NucBuster Protein Extraction Kit | Millipore | 71183 |
| MycoStrip Mycoplasma Detection Kit | InvivoGen | rep-mys-20 |
| Direct-zol RNA isolation kit | Zymo Research | R2052 |
| Mouse Cytokine/Chemokine 44-Plex Discovery Assay Array (assays performed by Eve Technologies) | Eve Technologies | MD44 |
| Precellys CK28-R Protein Safe Hard tissue homogenizing tubes | Bertin Technologies | P000972-LYSK0-A.0 |
| U-PLEX Custom Biomarker Group 1 Assay (Mouse) | Mesoscale Discovery | K15069M |
| **Experimental models (cell lines/mice)** | | |
| 293-FT (derived from human embryonal kidney cells transformed with the SV40 large T antigen) | Thermofisher Scientific | R70007 |
| Murine B16-F10 Cas9-expressing control | This study | Parental line is CRL-6475 from ATCC |
| Murine B16-F10 ΔUNG | This study | |
| Murine B16-F10 ΔUNG_2 | This study | |
| C57BL/6J mice | Jackson Laboratories | JAX:000664 |
| Athymic nude mice (NU/J) mice | Jackson Laboratories | JAX:002019 |

**Software and algorithms**

| Resource | Source | Link |
|---|---|---|
| StepOnePlus Real-Time PCR System | ThermoFisher Scientific | |
| GraphPad Prism 10.1.1 | GraphPad | https://www.graphpad.com/scientific-software/prism/ |
| FlowJo v10 | BD Biosciences | https://www.flowjo.com/solutions/flowjo |
| Gene set enrichment analysis (GSEA) 4.3.2 | | www.gsea-msigdb.org |
| ClusterProfiler | | https://bioconductor.org/packages/release/bioc/html/clusterProfiler.html |
| Gene Ontology | | http://www.geneontology.org |
| MSD Discovery Workbench | Mesoscale Discovery | https://www.mesoscale.com/en/products_and_services/software |
| Adobe Illustrator 2023 | Adobe | https://www.adobe.com/products/illustrator.html |
| Adobe Photoshop 2023 | Adobe | https://www.adobe.com/products/photoshop.html |

#### Lead contact

**EXTENDED METHODS DETAILS**

### Lentivirus production and cell transduction

Virus production: 293-FT cells were used to prepare lentivirus expressing Cas9 and Cas9/sgRNA specific to murine *Ung*. Cells ($1x10^6$/dish) were seeded in a 60mm dish for overnight incubation. Packaging vectors for the third-generation system pMDLg/pRRE (Addgene, Cat# 12251), pRSV-Rev (Addgene, Cat# 12253), and pMD2.G (Addgene, Cat# 12259) and the shuttle vectors/transfer vectors (see Key Resources Table) were co-transfected into 293-FT cells using the TransIT-X2 Dynamic Delivery System (Cat# MIR 6003). Supernatant containing the lentivirus was collected after 48 hours, followed by filtration using 0.45 µM filters to remove cell debris and isolate the viral particles. The lentivirus particles were then further concentrated using the Lenti-X Concentrator (Takara Bio, Cat# 631231) according to the manufacturer's instructions.

Transduction: Target cells ($2\times10^5$/well) were seeded into a 6-well plate and cultured for 24 hours. Next, the media was replaced with 1 mL of complete media followed by dropwise addition of the lentiviral particle solution (1 mL). After overnight incubation in the incubator, lentivirus-containing media was replaced with complete media with puromycin (1.5 µg/ml) and kept under selection for 5 days. Media/puromycin was replaced every other day.

Generation of *Ung* knockout cells: After puromycin selection, the polyclonal population was seeded into 96-well plates as single cells using the serial dilution method. Once cell clones reached confluency, they were harvested. Genomic DNA was isolated (PureLink Genomic DNA Mini Kit, Invitrogen) and amplified using Primer Pair 1 (see Key Resources Table). Next, amplified DNA was purified using the GeneJET PCR purification kit (ThermoFisher). Purified DNA with Primer Pair 2 (see Key Resources Table) was submitted to Azenta Life Science (New Jersey, USA) for Sanger sequencing to validate the knockout clone.

### Cell lysate preparation for the DNA repair molecular beacon assay

Harvested cells were centrifuged at 500 x g at 4°C for 5 minutes. The supernatant was discarded and the nuclear lysate from the cell pellets was extracted using the NucBuster Protein Extraction Kit (Millipore) with all steps performed on ice. The protein content of the resulting nuclear protein extract was measured using a Nanodrop 2000 Spectrophotometer. All nuclear lysates were diluted to 2 µg/uL in preparation for the DNA Repair Molecular Beacon (DRMB) assay. BER buffer was used to dilute the nuclear lysates.

### DNA repair molecular beacon assay

The DRMB assay was performed as follows:

Buffer Preparation: The DRMB assay is run in a standard Base Excision Repair (BER) Reaction buffer that is comprised of 25mM HEPES-KOH, 150mM KCl, 0.5mM EDTA, 2% Glycerol, and 0.5mM DTT in autoclaved water. BER buffer was filtered using a 0.45 µm filter.

Beacon and Annealing: DNA repair molecular beacons were annealed prior to use in the beacon assay to ensure the integrity of a hairpin structure. Four DNA repair molecular beacons were used (Con2, dU/A, dU/G, and G/dU) and all beacons contain a 6-Fam fluorophore on the 5' end and a Dabcyl non-fluorescent quencher on the 3' end (see see Key Resources Table). The dU/A probe contains deoxyuridine opposite adenine. The dU/G probe contains deoxyuridine opposite guanine. The G/dU also contains a deoxyuridine opposite guanine, but in the reverse order. Con2 contains no such modifications and is used as a negative control. All beacons were diluted to 200 nM in BER Reaction buffer in light-blocking microcentrifuge tubes. After dilution, beacons were denatured by boiling for 3 minutes and were then left overnight to anneal into hairpin structures.

Assay: Samples were loaded into a MicroAmp Fast Optical 96-Well Reaction Plate on ice, with each reaction containing 15 μL of BER Reaction buffer, 5 μL of beacon, and 5 μL of lysate (at 2 μg/uL). Each experimental sample was assayed in quadruplicate for each of the four beacons. Three additional controls, in quadruplicate, were included: buffer only (25 μL of BER buffer), buffer + lysate (20 μL buffer + 5 μL lysate), and buffer + beacon (20μL buffer + 5μL beacon). All sample lysates had corresponding buffer + lysate wells, and all four beacons had corresponding buffer + beacon wells. Plates were sealed with optical adhesive film, lightly vortexed, centrifuged at 500 x g for 10 seconds, and assayed using the StepOnePlus Real-Time PCR System. The PCR consisted of two stages. First, samples are maintained at 37°C and fluorescence was measured every 20 seconds for 180 cycles. These fluorescence readings represent the experimental data. Second, the temperature increases in 5 increments to 60°C, 65°C, 70°C, 75°C, and 80°C in succession with fluorescence measured every 20 seconds for a total of 15 cycles at each temperate. The purpose of this stage is to determine the temperature of maximum fluorescence (Tmax) for a given lysate + beacon combination. Dividing the experimental fluorescence by the Tmax for each beacon normalizes the data and converts relative fluorescence to normalized fluorescence. Normalized fluorescence values allow inter-plate comparisons.

Analysis: Each experimental value had corresponding 'buffer + lysate' and the 'buffer + beacon' values subtracted from the measured value. The 'buffer only' value was then added. The baseline was then adjusted by subtracting cycle 15 from each experimental value (ignoring all values before cycle 15) yielding baseline adjusted values. Fluorescence in the second stage was then analysed to determine the Tmax temperature. The baseline adjusted values were then divided by the fluorescence at Tmax to yield normalized fluorescence values for each experimental sample, in quadruplicate. The four values were averaged and plotted against time.

**Flow cytometry**

For *in vitro* flow cytometry experiments, Cas9-expressing control, ΔUNG, and ΔUNG_2 B16 cells were treated with 1.0 or 5.0 ng/mL recombinant mouse IFN-α1 (BioLegend), 0.2 or 1.0 ng/mL recombinant mouse IFN-β1 (BioLegend), 0.1, 0.3, or 1.0 ng/mL recombinant mouse IFN-γ (Peprotech, dissolved according to the manufacturer's instructions), or equivalent volumes of 1x PBS, for 18 h. Interferon-treated cells were harvested using Accutase (Gibco) cell dissociation reagent (IFN-α/β experiments) or were gently scraped for collection (IFN-γ experiments), pelleted, resuspended in flow cytometry staining (FCS) buffer (Invitrogen), and aliquoted to 96-well round-bottom plates. Cells were blocked in 100 μL 5% normal mouse serum (Invitrogen) in 1x PBS for 10 min on ice, stained with MHC-I and PD-L1 or isotype control antibodies (all 1:100 in 100 μL FCS buffer) for 1 h on ice in the dark, and stained with efluor780 fixable viability dye (1:3000 in 100 μL 1x PBS) for 10 min on ice in the dark, with FSC buffer washes between staining steps. Cells were analyzed live or were fixed in 175 μL FluoroFix (BioLegend) for 2 h at room temperature in the dark and washed in FSC buffer prior to analyses. Single-stained OneComp eBeads (Invitrogen) were used as compensation controls for MHC-I and PD-L1. A single-stained mix of live and heat-killed (30 sec at 95°C) cells were used as the compensation control for efluor780. Acquisition was performed with a 4-laser CytoFLEX (Beckman Coulter) and analyses were performed in FlowJo V10.

For *in vivo* immune profiling experiments, Cas9-expressing control or ΔUNG B16 tumors were harvested from mice at the indicated time points. Corresponding splenocytes were used for single color controls, fluorescence-minus-one (FMO) controls, and for general gating. Tumors were weighed prior to processing. Single cell suspensions were generated from tumors and spleens. Tumor tissue (≤ 250 mg) was minced and then digested in Collagenase IV Cocktail (approximately 2 mL per 100 mg tumor) containing 3.2 mg/mL Collagenase IV, 1 mg/mL DNAse I, 2 mg/mL Soybean Trysin Inhibitor (all Worthington) for two 15-minute incubations at 37°C, with periodic agitation, and tituration steps between incubations and after the second incubation. Tumor homogenate was smashed through a 70 µm cell strainer (Corning) using the rubber plunger of a syringe and the filter was rinsed with 1x PBS. Tumor samples were centrifuged at 350 x g for 5 minutes and pellets were resuspending in complete D-MEM media. Spleens were mechanically dissociated between frosted glass slides and filtered through 70 µm cell strainers. Erythrocytes were lysed in 150 mM NH4Cl, 10 mM NaHCO3, 0.1 mM EDTA pH 8.0. for 10 sec (tumors) or 30 sec (spleens). Cell suspensions were counted with a Scepter 3.0 (Millipore) and seeded at 1.2-2 x $10^6$ cells (equivalent number within an experiment) in 96-well round bottom plates for blocking and staining as follows: Fc receptors were blocked for 10 min at 4°C with 0.5 µg anti-CD16/32 antibody (TruStain FcX Plus, BioLegend) in FSC buffer (Invitrogen), cells were stained with antibodies to surface antigens (in FSC buffer) for 15 min at 4°C, dead/dying cells were stained with eFlour780 fixable viability dye (1:2000-3000, Invitrogen) or LIVE/DEAD fixable near-IR dead cell stain (1:1000, Invitrogen) in 1x PBS for 10 minutes at 4°C, samples were fixed and permeabilized in eBioscience Fixation/Permeabilization reagent (Invitrogen) for 15 min at room temperature, and when performing nuclear (Ki67, Foxp3) or intracellular cytokine (IFN-γ, TNF-α, IL-17) staining, samples were stained for 45 min at room temperature in eBioscience 1x Permeabilization Buffer (Invitrogen) containing antibodies to nuclear/intracellular proteins. Brilliant Stain Buffer Plus (BD Biosciences) was added to antibody cocktails containing multiple Brilliant Violet dye conjugates to prevent polymer dye-dye interactions. True-Stain Monocyte Blocker (BioLegend) was added to surface antibody cocktails containing PE-Cy7 conjugates to prevent non-specific binding of monocytes/macrophages to the tandem dye. For measurement of cytokine-producing $CD8^+$ and $CD4^+$ T cells, prior to staining, cells were stimulated for 4 h with 1x eBioscience Cell Stimulation Cocktail (plus protein transport inhibitors) (Invitrogen), which contains PMA and ionomycin, in complete D-MEM media. Unstimulated controls were treated for 4 h with 1x eBioscience Cell Protein Transport Inhibitors cocktail (Invitrogen) in D-MEM media. Since PMA/ionomycin stimulation induces internalization of CD3 and the CD8/CD4 co-receptors, staining for CD3, CD4, CD8, and CD45 was performed post-fixation/permeabilization during intracellular cytokine staining. Data were acquired uncompensated data using a BD LSRFortessa 4-laser cytometer and BD FACSDiva software, with compensation performed in FlowJo V10 software, or data were acquired compensated/unmixed using a Cytek Aurora spectral cytometer and SpectroFlow software. Single stained spleen samples with matching unstained cells or single stained OneComp eBeads or UltraComp eBeads Plus (Invitrogen) were used for single color compensation controls. Fluorescence-minus-one (FMO) controls were used, where appropriate, to empirically determine gating. All data analyses were performed in FlowJo V10 software. Gating strategies are shown in Supplemental Figures S8-S12.

**RNA extraction and RNA sequencing**

Cells were seeded in 35 mm dishes and cultured overnight. The next day, cells were treated with 5 μM ATRi AZD6738 for 48 h. At the time of harvest, cells were washed once with 1x PBS, resuspended in 1x PBS, and Trizol (in a 1:3 ratio) was added to the cellular suspension. Total RNA was extracted using the Direct Zol RNA kit (Zymo Research Corp, Irvine) according to the manufacturer's instructions. The isolated RNA from two biological replicates per condition was submitted to the Novogen Advancing Genomics facility (Sacramento, CA, USA) for RNA sequencing. For RNA-seq analyses, the original data file from a high-throughput sequence platform was transferred to sequence read by CASAVA base recognition and stored in FASTQ format files. Raw data were further processed by removing adapter reads and reads with uncertain nucleotide constituting more than 10%. Read alignment was performed by Novogen using HISAT2 software and the mouse reference genome. FPKM (Fragments per kilobase of transcripts sequence per millions base pairs sequenced) was used to quantify the abundance of transcripts or genes. FPKM values and the number of raw and clean reads for each sample, provided by Novogen, are included in the supplemental spreadsheet. Differential gene expression analysis was performed using DEseq2 (for biological replicates) or edgeR (for no biological replicates) software with FDR correction by the Benjamini-Hochberg procedure with threshold [log2(foldchange)]>=1 & p-adj <=0.05. Differential gene expression analysis was performed by Novogen. Gene set enrichment analysis (GSEA 4.3.2) was utilized to perform rank-based identification of the most enriched pathways between groups. One biological replicate for ATRi-treated ΔUNG_2 was excluded from GSEA in Figure 3 because the raw data (BAM) file was corrupt and could not be deposited to Gene Expression Omnibus (GEO). The analyzed data for this replicate, provided by Novogen, are included in the supplemental spreadsheet for reference (sample CB_10). Group pathway analysis was performed using ClusterProfiler software for Gene Ontology (GO) analysis (http://www.geneontology.org/). GO terms with p-adj <0.05 are significantly enriched, and the most significant terms were selected for display.

**REFERENCES**


1. Li J, Svilar D, McClellan S, Kim JH, Ahn EE, Vens C, et al. DNA Repair Molecular Beacon assay: a platform for real-time functional analysis of cellular DNA repair capacity. *Oncotarget.* 2018;9(60):31719-43.
2. Svilar D, Vens C, and Sobol RW. Quantitative, real-time analysis of base excision repair activity in cell lysates utilizing lesion-specific molecular beacons. *J Vis Exp.* 2012(66):e4168.
